# Structure-aware *M. tuberculosis* functional annotation uncloaks resistance, metabolic, and virulence genes

**DOI:** 10.1101/358986

**Authors:** Samuel J Modlin, Afif Elghraoui, Deepika Gunasekaran, Alyssa M Zlotnicki, Nicholas A Dillon, Nermeeta Dhillon, Norman Kuo, Cassidy Robinhold, Carmela K Chan, Anthony D Baughn, Faramarz Valafar

## Abstract

Accurate and timely functional genome annotation is essential for translating basic pathogen research into clinically impactful advances. Here, through literature curation and structure-function inference, we systematically update the functional genome annotation of *Mycobacterium tuberculosis* virulent type strain H37Rv. First, we systematically curated annotations for 589 genes from 662 publications, including 282 gene products absent from leading databases. Second, we modeled 1,711 under-annotated proteins and developed a semi-automated pipeline that captured shared function between 400 protein models and structural matches of known function on protein data bank, including drug efflux proteins, metabolic enzymes, and virulence factors. In aggregate, these structure- and literature-derived annotations update 940/1,725 under-annotated H37Rv genes and generate hundreds of functional hypotheses. Retrospectively applying the annotation to a recent whole-genome transposon mutant screen provided missing function for 48% (13/27) of under-annotated genes altering antibiotic efficacy and 33% (23/69) required for persistence during mouse TB infection. Prospective application of the protein models enabled us to functionally interpret novel laboratory generated Pyrazinamide-resistant (PZA) mutants of unknown function, which implicated the emerging Coenzyme A depletion model of PZA action in the mutants’ PZA resistance. Our findings demonstrate the functional insight gained by integrating structural modeling and systematic literature curation, even for widely studied microorganisms. Functional annotations and protein structure models are available at https://tuberculosis.sdsu.edu/H37Rv in human- and machine-readable formats.

**IMPORTANCE:** *Mycobacterium tuberculosis*, the primary causative agent of tuberculosis, kills more humans than any other infectious bacteria. Yet 40% of its genome is functionally uncharacterized, leaving much about the genetic basis of its resistance to antibiotics, capacity to withstand host immunity, and basic metabolism yet undiscovered. Irregular literature curation for functional annotation contributes to this gap. We systematically curated functions from literature and structural similarity for over half of poorly characterized genes, expanding the functionally annotated *Mycobacterium tuberculosis* proteome. Applying this updated annotation to recent *in vivo* functional screens added functional information to dozens of clinically pertinent proteins described as having unknown function. Integrating the annotations with a prospective functional screen identified new mutants resistant to a first-line TB drug supporting an emerging hypothesis for its mode of action. These improvements in functional interpretation of clinically informative studies underscores the translational value of this functional knowledge. Structure-derived annotations identify hundreds of high-confidence candidates for mechanisms of antibiotic resistance, virulence factors, and basic metabolism; other functions key in clinical and basic tuberculosis research. More broadly, it provides a systematic framework for improving prokaryotic reference annotations.

## INTRODUCTION

Manual curation remains the gold standard for annotating function from literature(1), yet requires massive effort from highly specialized researchers. UniProt curators alone evaluate over 4,500 papers each year(1). Literature annotation is typically complemented with functional inference by sequence homology, but this approach fails to identify distant relatives (remote homologs) or convergently evolved proteins of shared function (structural analogs).

These challenges hinder the study of *Mycobacterium tuberculosis*, the etiological agent of tuberculosis. The *M. tuberculosis* virulent type strain is H37Rv, a descendant of strain H37, was isolated from a pulmonary TB patient in 1905 and kept viable through repeated subculturing(2). Following sequencing of the H37Rv genome, function was assigned to 40% of its open reading frames (ORFs)(3), then expanded to 52% in 2002 following re-annotation(4). New annotations continued to be added by TubercuList (now part of Mycobrowser, https://mycobrowser.epfl.ch/) until March 2013. To date, one-quarter of the H37Rv genome (1,057 genes) lacks annotation entirely, listed in “conserved hypotheticals” or “unknown” functional categories and hundreds more minimally describe product function (e.g. “possible membrane protein”). Though other databases have emerged in recent years(5-9) Mycobrowser remains the primary resource for gene annotation for TB researchers(10) yet lacks functional characterizations reported in the literature.

Moreover, many proteins key to *M. tuberculosis* pathogenesis are challenging to ascribe function by sequence similarity. For instance, transport proteins—many of which allow *M. tuberculosis* to tolerate drug exposure by effluxing drug out of the cell(11)—have membrane-embedded regions under relaxed constraint compared to globular proteins and diverge in sequence more rapidly as a result(12). This rapid divergence challenges their characterization through homology. Limitations of sequence-based approaches to detect and annotate *M. tuberculosis* proteins motivates an alternative approach to annotating *M. tuberculosis* gene function.

One alternative approach is identifying protein homologs and analogs through shared structure, which offers considerable advantages. First, it removes bias toward *a priori* assumptions by not limiting search space to evolutionarily close relatives, facilitating discovery of functions unexpected by conventional predictions. Second, structure-based functional annotation can infer analogy between proteins structures. This ability to detect analogy is especially valuable for inferring function at the host-pathogen interface, which is challenging to recapitulate in the laboratory. Moreover, analogous relationships between proteins of shared structure/function cannot be resolved by sequence homology because they often evolve convergently with low amino acid (AA) similarity rather than from common ancestry (13).

Iterative Threading ASSEmbly Refinement (I-TASSER)(14), builds three-dimensional protein structure from sequence through multiple threading alignment of Protein Data Bank (PDB)(15) templates, followed by iterative fragment assembly simulations. I-TASSER accurately predicts structure (16-20), provides metrics for model quality(21) (C-score) and pairwise structural similarity(22) (TM-score), and integrates function and structure prediction tools(23) (COACH and COFACTOR) comprising Gene Ontology (GO) terms(24), Enzyme Commission (EC) numbers(25), and Ligand Binding Sites (LBS)(26).

EC numbers and GO terms partially or completely define gene function and are widely incorporated into mainstream databases. EC numbers describe catalytic function hierarchically through a four-tiered numerical identifier system that funnels from general enzyme class (e.g. ligase, oxidoreductase) down to substrate specificity with atomic precision(25). GO terms add to EC number content: they describe gene products by where they function, the processes they are involved in, and their specific molecular function in species-independent form(27, 28). This cross-species unification is particularly useful for reconciling annotation transfers of analogs and distant homologs into gene product names.

Previous hypothetical gene annotation efforts for *M. tuberculosis* have not included a systematic manual literature curation component and have drawn from inferential techniques such as protein homology and fold similarity(29, 30), aggregating gene orthology server predictions(31), metabolic pathway gap-filling(32), and STRING interactions(33), lacking inclusion criteria based on benchmarked likelihood of correctness. Measured interpretation of annotated gene functions requires the source of the annotation and the reliability of the evidence warranting it to be described explicitly. We strived to provide this resource by reconciling the H37Rv annotation on Mycobrowser with published functional characterization and systematically inferring function from structural similarity to annotate genes challenging to characterize through experiment and sequence analysis. We include orthogonal validation measures to confidently capture unexpected functions while minimizing “overannotation”(34, 35).

We report our findings in three sections. First, we establish the set of under-annotated genes, describe our systematic manual literature curation protocol, and summarize the novelty of the resulting annotations with respect to popular functional databases. Next, we describe our structural modelling pipeline, orthogonal validation and quality assurance methods, and two illustrative examples of manually curated functional annotations from structural inferences unsupported by an established method of detecting remote functional homology. Finally, we summarize the updated annotation and genes remaining to be characterized, and demonstrate the value added of this annotation through its application to previously published and novel functional screens.

## RESULTS

### Numerous genes lack annotation in all common *M. tuberculosis* databases

First, we defined a set of 1,725 under-annotated genes (Dataset 1) based on their TubercuList entry. We included:

1. Genes in “conserved hypothetical” or “unknown” functional categories.
2. Genes qualified by an adjective connoting low confidence (e.g. “predicted” or “possible”).
3. Genes described by something other than function: (e.g. “Alanine-rich protein” or “isoniazid-inducible protein”).
4. PE/PPE family genes – A largely uncharacterized, polymorphic protein family unique to Mycobacteria with proline-glutamine or proline-proline-glutamine N-terminal domains.

Next, we asked how many of these genes lacked annotations across commonly referenced databases (Table 1). Although BioCyc and UniProt had more genes with GO terms than TubercuList, and UniProt, and Mtb Network Portal had fewer hypothetical proteins than TubercuList, all databases had over one-quarter of coding sequences (CDS) annotated as hypothetical, demonstrating the need for systematic manual annotation.

**Table 1.**
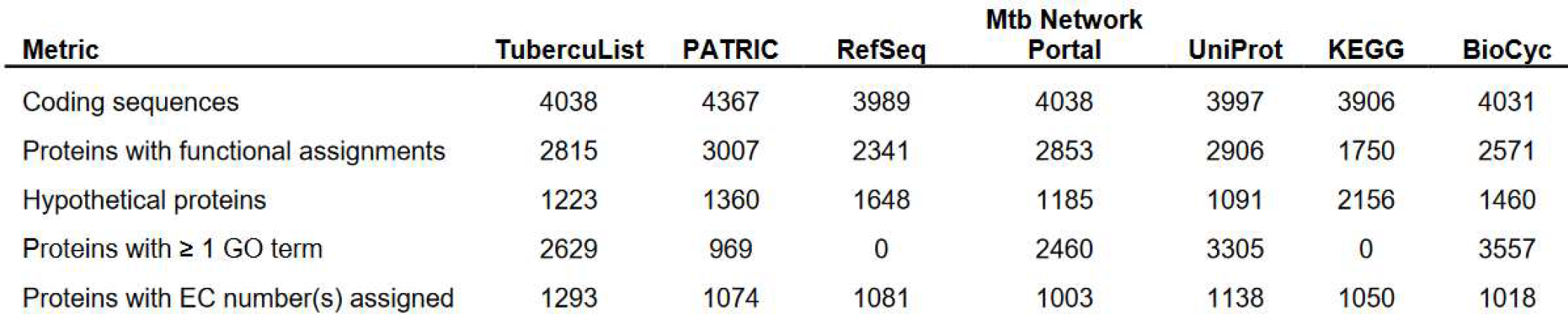
Comparison among frequented annotation resources. “Functional assignments” refer to annotations that describe protein function, and excluded hypothetical, unknown/uncharacterized, and PE/PPE family proteins. Counts reflect database content on May 17, 2017 for RefSeq(36) https://www.ncbi.nlm.nih.gov/refseq/, PATRIC(6) https://www.patricbrc.org/,and Mtb Network Portal(9) http://networks.systemsbiology.net/mtb/ and June 23, 2017 for KEGG(120) https://www.kegg.jp/kegg/genome/pathogen.html and UniProt(116) https://www.uniprot.org/uniprot/. The number of CDSs in KEGG is reported as 3906 because they include only protein coding genes. The source of annotations for *M. tuberculosis* protein-coding genes in KEGG is TubercuList(125).

### Frequently consulted annotation sources lack experimentally demonstrated functions

We devised a manual curation protocol (*Text S1*, Fig. S1) that:

1. Assigns qualifying adjectives that connote confidence
2. Assigns Enzyme Commission (“EC”) numbers
3. Requires multiple reviewers per paper to hedge against human error, and an additional quality control curator to check formatting and annotation consistency.

We systematically reviewed over 5,000 publications according to this protocol, furnishing annotations for one-third of under-annotated genes (575) with product function or functional notes (Dataset 1). Of these, 282 were annotated with product function absent from TubercuList, including 122 enzymes and 28 regulatory proteins. These annotations include fourteen oxidative stress response genes, 22 proteins mediating RNA and DNA functions, and eight transport/efflux proteins.

Next, we evaluated whether these missing annotations were restricted to TubercuList or more widespread. We checked our curations against four frequented annotation resources: UniProt (Dataset 2), Mtb Network Portal(9), PATRIC(6), and RefSeq(36) (Dataset 2). Product function information was absent from 172/282 (61%) of these genes on UniProt (Dataset 2) and 118 (64 of which are antigens, *Text S1*, Fig. S4A) were more thoroughly annotated than in any of the examined databases (Dataset 2). This novelty underscores the value of these manual curations and highlights critical information that these databases lack (Table 2 and Dataset 1). After excluding antigens, 25.2% of genes with function curated from literature were absent from all five annotation resources. To identify enzymatic functions unannotated elsewhere, we compared our manual EC number assignments to commonly referenced databases (*Text S1*, Fig. S4). This comparison revealed 59/98 (60.2%) of genes assigned EC numbers have EC numbers only in our annotation. These missing annotations include functions affecting drug-resistance, features of *in vivo* infection, and other important functions. Examples include a rare instance where a PE/PPE gene has demonstrated catalytic function(37) (Rv1430), a probable peptidoglycan hydrolase implicated in isoniazid (INH) and PZA resistance and biofilm formation(38) (Rv0024), a rhomboid protease with roles in biofilm formation and ciprofloxacin and novobiocin resistance(39) (Rv1337), and an oxidoreductase important for in-host survival of *M. tuberculosis*(40) (41, 42) (Rv3005c). Additional findings pertinent to pathogenesis, host-pathogen interaction, and antibiotic resistance were noted across under-annotated genes (S1 Annotation).

**Table 2.**
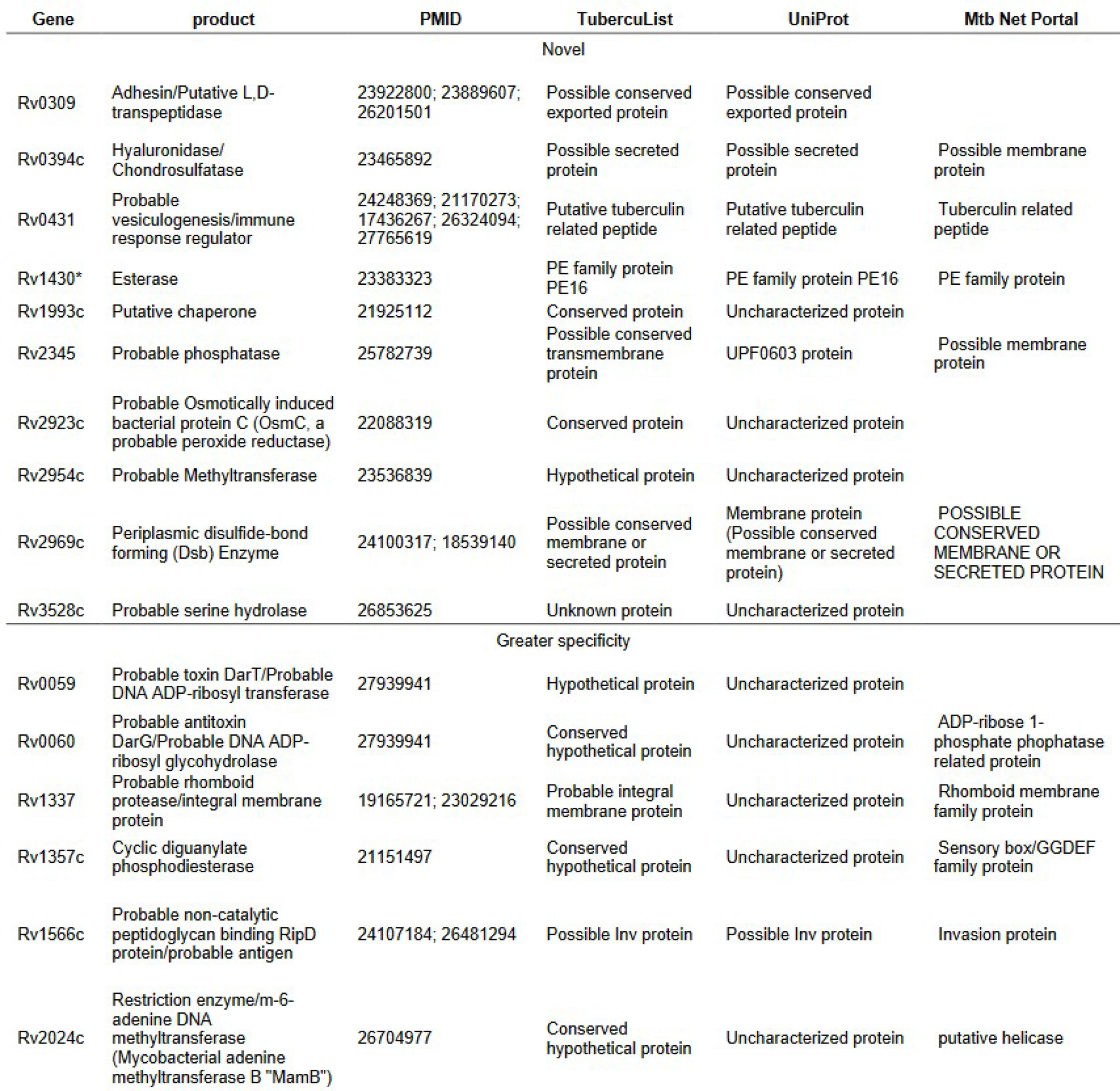
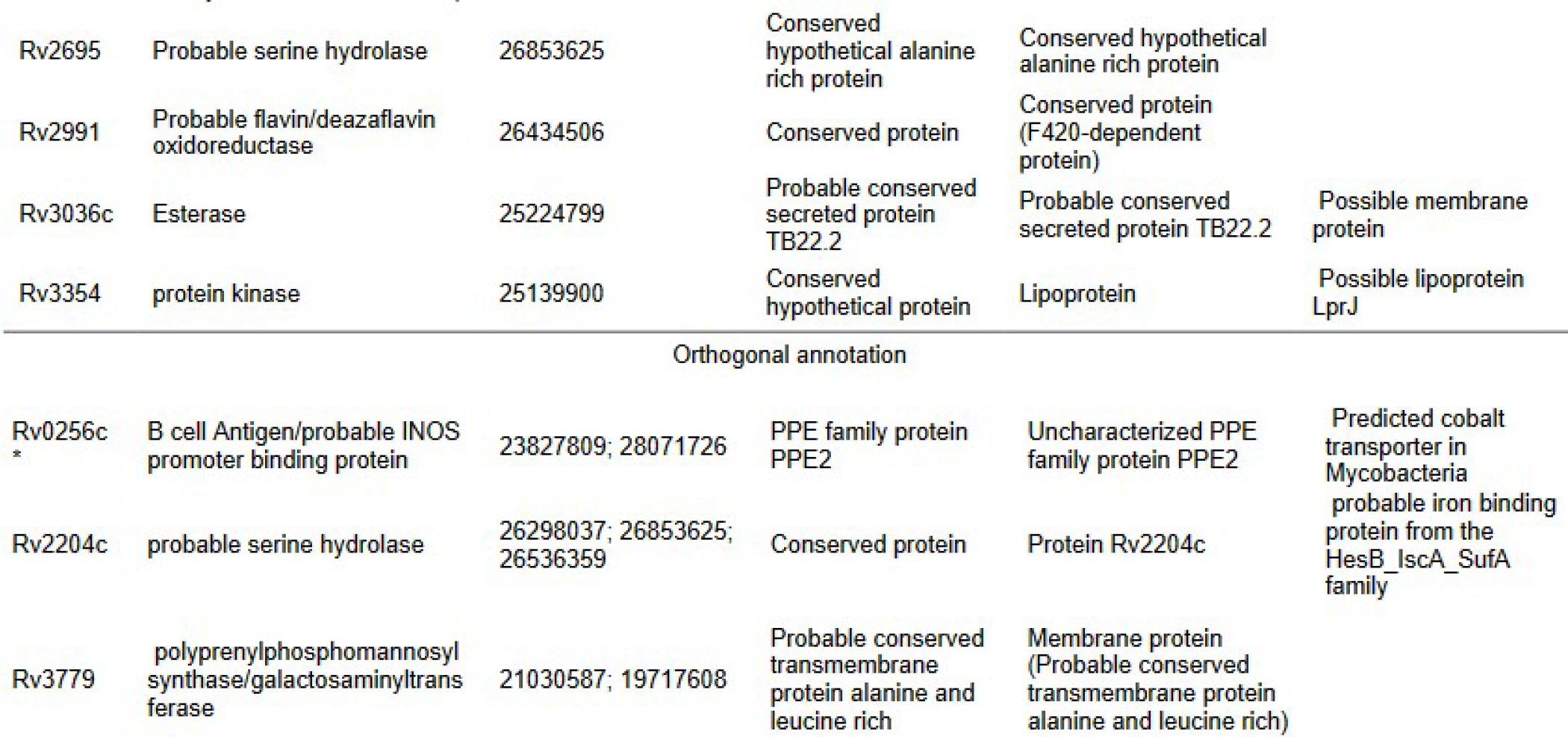
Genes annotated through systematic manual curation that expand upon annotations from major databases. Annotations are separated into completely novel, those with similar annotations but with greater specificity, and those with an additional, orthogonal annotation compared to evaluated databases (*Text S1*, Fig. S4A). Pubmed IDs (PMID) from which annotations for each product were derived are included. Members of the PE/PPE family are indicated by asterisk.

### Annotating function from structure similarity

Next, we modeled protein structures and developed a procedure to annotate function based on shared structure according to the likelihood that two proteins shared function (i.e. precision; Equation 1, Methods and Materials).

To inform our annotation methods we first assessed whether we could

1. Reliably infer precision according to similarity.
2. Differentiate between precision thresholds at different levels of functional detail (e.g. EC number tiers).

To make these assessments we benchmarked precision as a function of template-modelling score (“TM-score”), a measure of structural similarity independent of protein length, and sequence similarity (Amino acid identity, “AA%”), using a set of 363 genes with known function (Methods & Materials) through the standalone version of I-TASSER. TM-score and AA% were predictive of precision and mutually correlated (*R*=0.784, Pearson correlation coefficient) among both concordant and discordant EC numbers (*Text S1*, Fig. S2B). We accounted for TM-score and AA% simultaneously by their geometric mean (µ_geom_) to estimate precision. Precision of EC number prediction increased monotonically as a function of µ_geom_ for all 4 EC tiers, and regression lines for the 4 degrees of EC functional specificity did not intersect (Fig. 1).

**Fig. 1.**
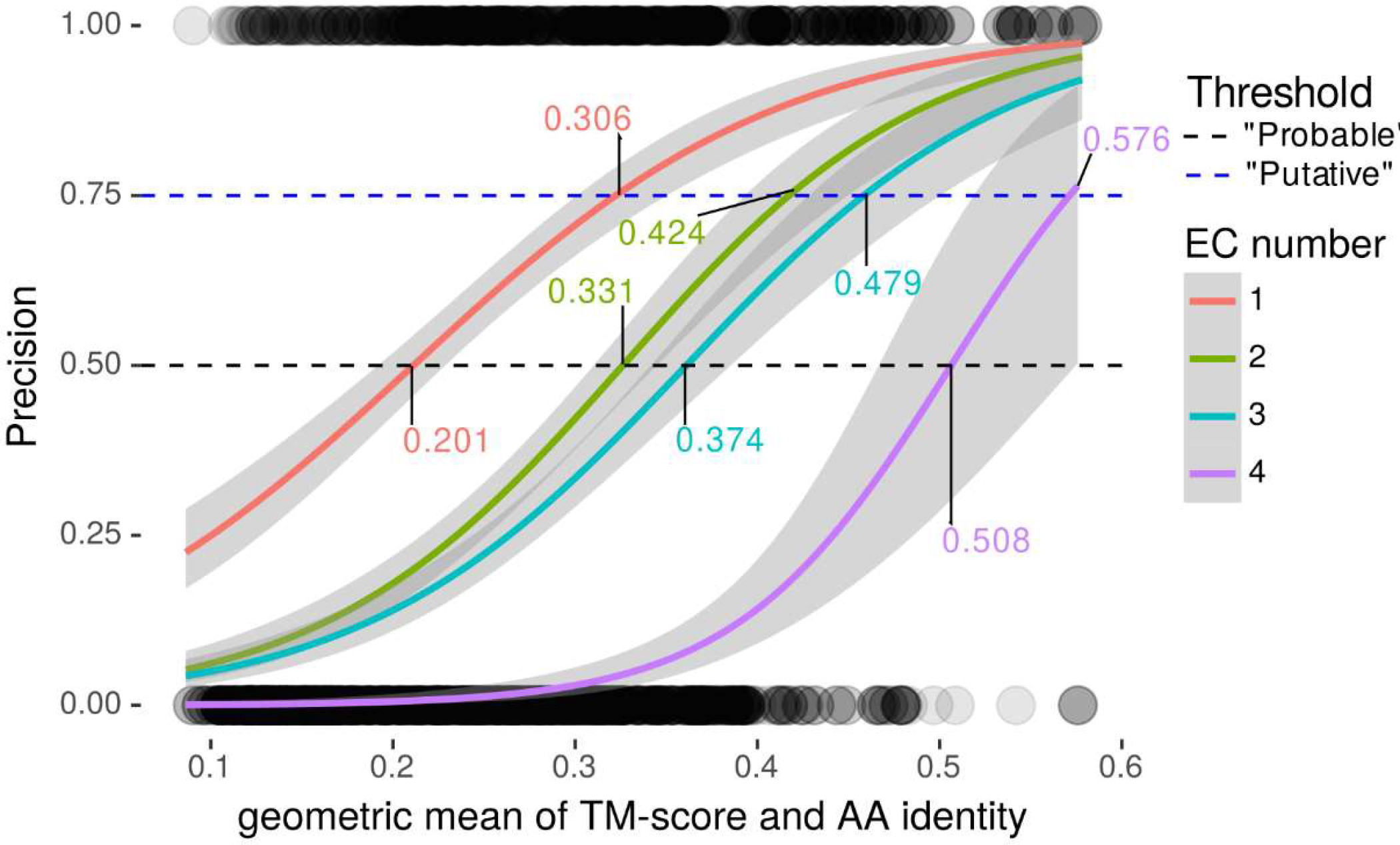
Determining similarity thresholds for annotation inclusion criteria. Precision of EC number as regressed against the geometric mean of TM-score and AA% (“µ_geom_”) for each specificity tier. Horizontal lines define (50% and 75%) thresholds, the points where precision intersects with regression lines for each EC specificity curve (labelled). Circles at the bottom and top are individual data points (incorrect = 0 and correct = 1, y-axis: precision, x-axis: µ_geom_). Circles are rendered at 10% opacity to visualize observation density. Only templates with AA% < 40% were included.

From these properties we concluded that we could reliably estimate precision from µ_geom_ with distinct thresholds for each EC tier. We defined thresholds as the µ_geom_ value where logistic regression lines intersected with 50% for receiving “putative” and 75% for receiving “probable” as qualifying adjectives (Fig. 1). Through this procedure we defined distinct thresholds for ascribing “putative” or “probable” status to enzymatic function at each of the 4 tiers of EC specificity. We incorporated EC numbers and GO terms from similar structures deposited on Protein Data Bank (PDB) hierarchically, according to evidence reliability (*Text S1*, Fig. S3). After the quality control pipeline described below, we recorded annotations in NCBI Table File Format and according to Genbank Prokaryotic Annotation Guide (www.ncbi.nlm.nih.gov/genbank/genomesubmit_annotation/) syntax and guidelines (integrated with manual curations from literature), and collated them into a unified functional annotation in GFF3 format (Fig. 2).

**Fig. 2.**
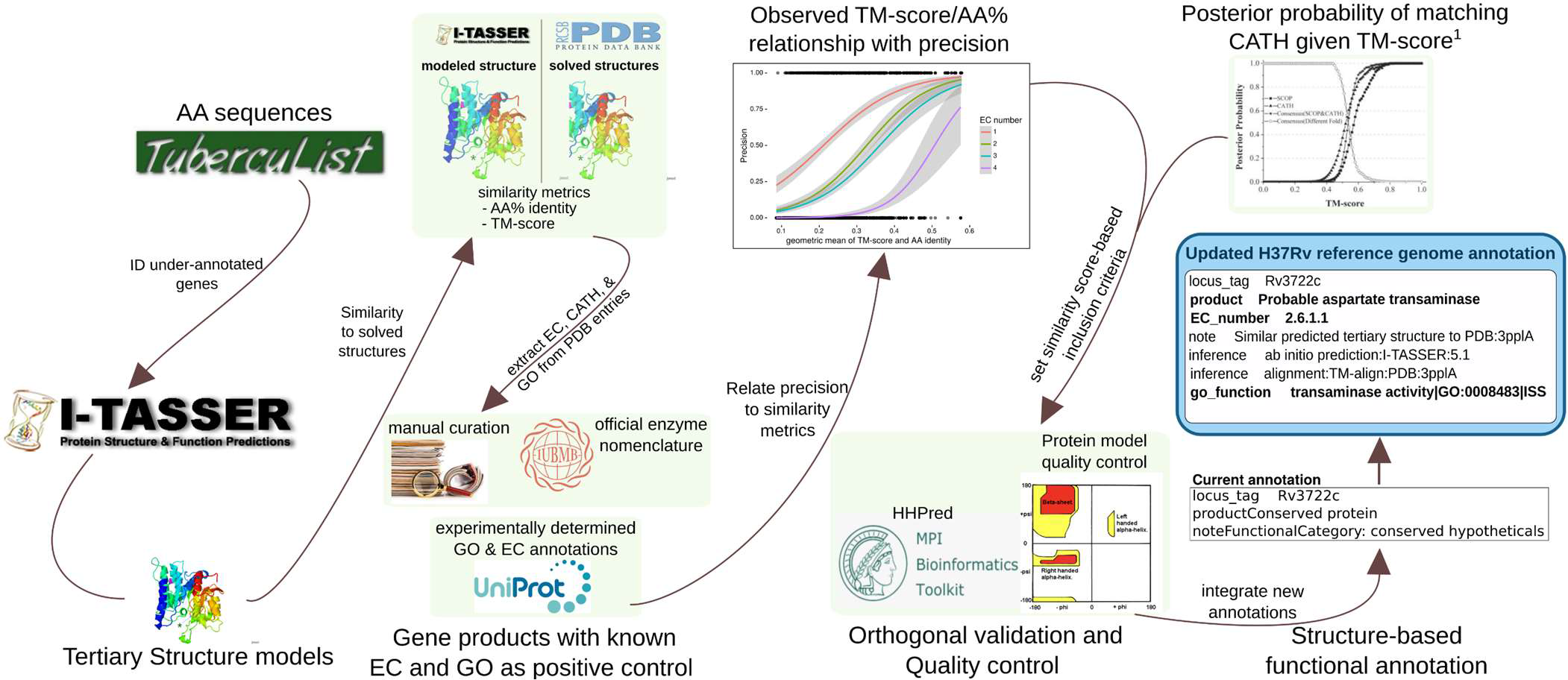
Information flow for producing annotations from structural similarity. The flow of information and procedures for acquiring, processing, filtering, and representing information, running from retrieval of amino acid sequences to the final updated H37Rv annotation. Some details are omitted for clarity. The 1,725 amino acid sequences were retrieved from TubercuList and run through a local installation of I-TASSER v5.1. 1,711 of 1,725 Amino Acid sequences had models generated successfully. Comparison metrics for sequence (Amino acid identity) and Structure (TM-score) were extracted from I-TASSER output. To set criteria for annotation transfer, precision (Equation 1) of GO Term and EC number concordance between similar matches on PDB and true function of 363 positive controls with GO terms and EC numbers of known function were regressed against extracted similarity metrics to generate a curve relating the geometric mean of TM-score and Amino acid similarity to precision. These informed inclusion thresholds for transferring GO and EC annotations from structures on PDB similar to the 1,711 modeled structures. CATH Topology folds were transferred according to a previous precision curve based on TM-score. This threshold was also used for inclusion of proteins classes that vary in sequence more than structure (e.g. transporters) and as criteria for transferring annotations from structures that were not annotated with EC numbers or GO terms. Annotations derived only from structure were passed through orthogonal validation and manual structure analysis for verification that transferred annotations were reasonable. All annotations were programmatically collated into an updated H37Rv reference genome annotation.

Although using µ_geom_ to determine inclusion criteria is useful for proteins with PDB entries of somewhat homologous sequence, it would not capture relationships by structural analogy or remote homology (because their low AA% would lower their score). To identify potential analogs and remote homologs, we used “TM_ADJ_”, an adjusted TM-score that accounts for model quality to conservatively estimate the TM-score between the true structure of a modelled protein and its putative homolog/analog of solved structure (Materials & Methods). We re-examined hits with TM_ADJ_ values that indicated matching topology according to previous benchmarks(21) (*Text S1*), and annotated function with EC numbers, GO terms, and product names (*Text S1* and Methods and Materials).

### Validating structure-based annotations

To validate our structure-based functional inference approach, we ran proteins with annotations derived only from structural similarity (n=366 through HHpred (43) (Fig. 3), a server that detects remote homology between proteins by comparing hidden Markov model profiles(43). We compared enzymatic structure-derived annotations (those with EC numbers, n=335 distinct EC number annotations from 271 proteins) programmatically, and non-enzymatic annotations manually (n=95, Dataset 3, Materials & Methods). Evaluating only the annotations to at least the 2^nd^ EC number level (n=325), most structure-inferred predictions were partially (288/335, 86.0%) or wholly (266/335, 79.4%) corroborated by HHpred (Fig. 3C), substantiating the validity of our structure-based approach to functional inference. Partially corroborated annotations (e.g. 3.1.2.4 to the level of 3.1.2.-, but not the 4^th^ level of EC specificity) were revised to reflect the less specific, HHpred-supported level of functional detail, and manually reconciled in cases where multiple EC numbers were corroborated (Materials & Methods).

**Fig. 3.**
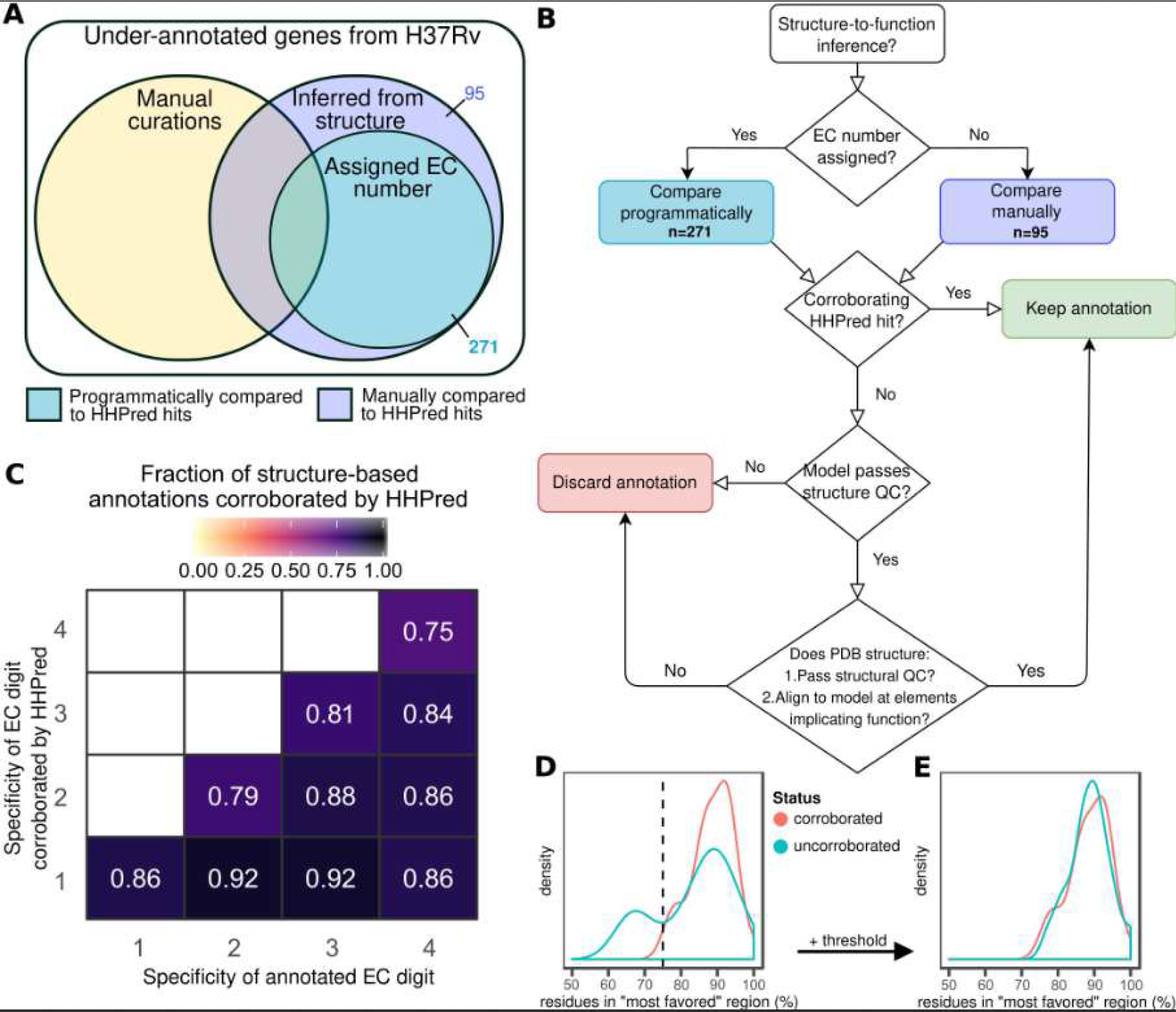
Orthogonal validation and quality assurance for structure-to-function inference. (**A**) Quality assurance and validation protocol assignment and (**B**) Decision workflow for retaining functional annotations inferred from structural similarity to proteins of solved structure and known function. (**C**) Heat map depicting fraction of EC number inferences corroborated by HHpred at each level of specificity. Fraction denominator is binned according to the number of E.C digits annotated (x-axis). (**D,E**) Structure quality assurance. Distribution of fraction of residues in “most favorable” region of Ramachandran plot prior to (**D**) and following application of a heuristic threshold (**E**) to discard biophysically improbable structural models.

Next, we assessed protein structure model quality using the fraction of residues in “most favored” regions of Ramachandran plots (Materials & Methods). Screening for abnormally low fractions can identify models with sterically untenable residue configurations, signaling low model quality(44). A threshold of 90% is often used for solved proteins(45), but we expected deviation from 90% even in quality models (since they are models rather than solved structures). To determine an acceptable threshold, we compared the distribution of residue fractions in “most favorable” regions among models with functions fully corroborated by HHpred with that of 29 wholly uncorroborated by HHpred. Fractions for HHpred-corroborated proteins distributed unimodally and peaked around 90% of residues falling in the “most favorable” region (median = 89.15%). This observation is consistent with HHpred-corroborated proteins having high-quality structures and informs us of the range of fractions characteristic of high-quality structural models. Models with functions wholly uncorroborated by HHpred, meanwhile, distributed bimodally, with one mode resembling the fully corroborated distribution and the second mode peaking at a lower fraction (Fig. 3D). This bimodal distribution is consistent with a mixture of quality models and truly poor models. To distinguish between poor- and high-quality models in the wholly uncorroborated set, we implemented a heuristic threshold at the intersection of the two distributions (75%, Fig. 3D). After removing models below the threshold, the remaining uncorroborated structures formed a single peak that resembled the HHpred-corroborated proteins (Fig. 3E). We used this threshold (75%) as the minimum acceptable fraction for HHpred-uncorroborated proteins to be considered for structure-based functional annotation.

Seven of the protein models with exclusively wholly unsupported structure-based annotations (n=29) were PE_PGRS protein models that resembled fatty acid synthase (FAS) subunit protein structures (particularly *Saccharomyces cerevisiae* PDB template 2pff). All seven failed Ramachandran filtering. This underscores the importance of these QC steps and suggests they excluded models implicating false functional analogies as intended. These annotations were likely artifactual, owing to glycine-abundant, low complexity regions of PE_PGRS proteins aligning to the hydrophobic regions of large eukaryotic synthases, inflating their similarity score and spuriously implying structural similarity.

Since HHpred is designed to detect homology between proteins(43) (but not necessarily analogy), there may be genuine functions inferred by our structural similarity pipeline that HHpred did not corroborate. To preserve such annotations while ensuring annotation quality, we manually inspected HHpred-uncorroborated annotations (Fig. 3B) for protein models that passed Ramachandran filtering (n=22). To accept annotations, we verified template protein quality, structural alignment of regions underlying function, and conservation of structural features and key functional residues. This step salvaged structure-derived functional annotations for nine proteins (Table 3, Dataset 3), two of which we highlight in detail in Fig. 4.

**Fig. 4.**
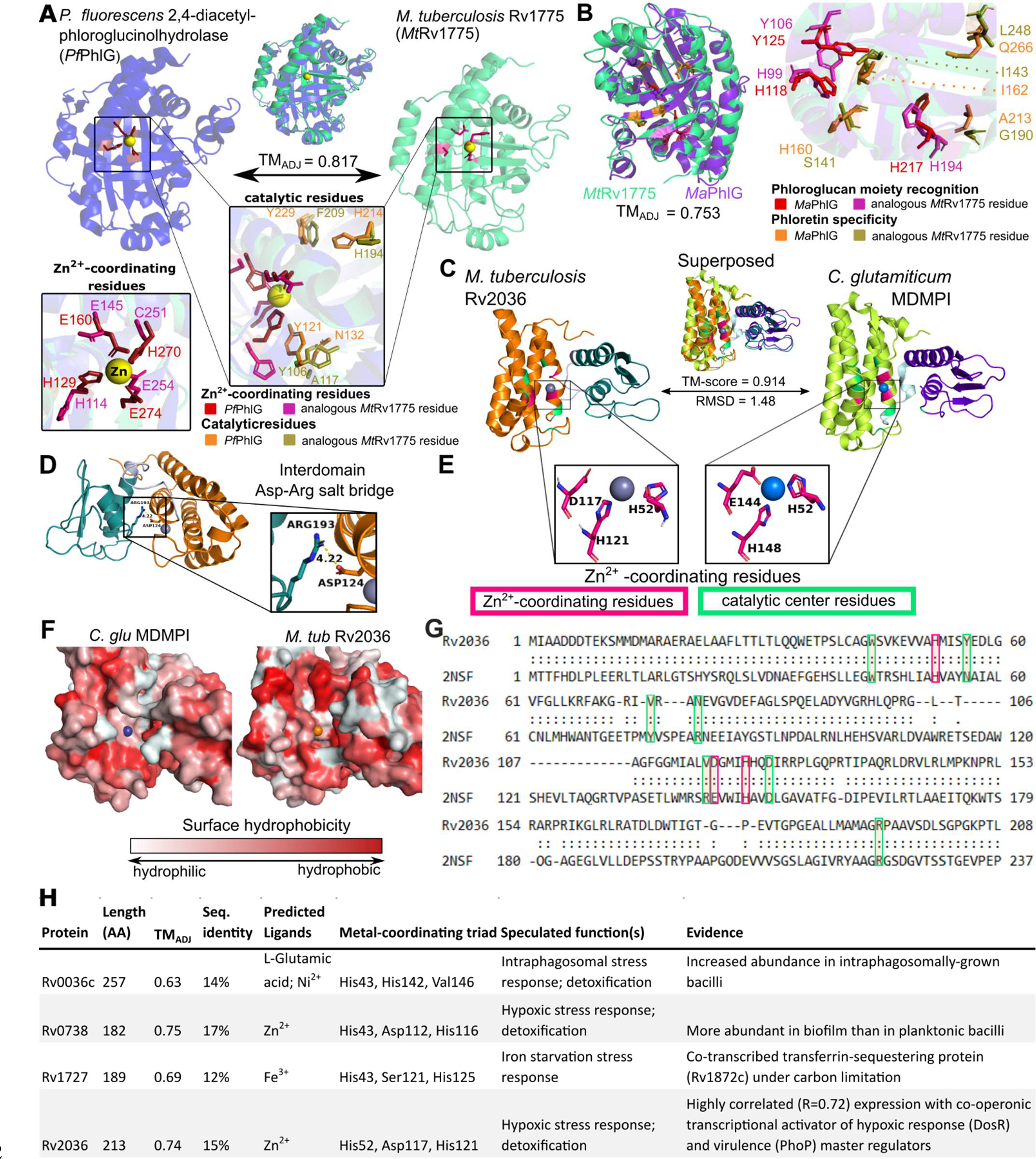
Manual structural analysis refines functional annotations uncorroborated by HHpred. **(A-B)** Conservation of structure- and sequence-features essential for C—C bond hydrolysis supports the inferred hydrolase function of Rv1775. **(A)** Structural alignment of modeled Rv1775 and its closest structural match (PDB 3hpw), a 2,4-diacetyl-phloroglucinolhydrolase of *Pseudomonas fluorescens* (*Pf*PhlG), The structures are superposed (top). Zoomed and re-oriented images of *Pf*PhlG Zinc-coordinating (box on left) and catalytic (pop-out) residues superposed with analogous *Mt*Rv1775 residues. **(B)** Comparison of functional and structural features between *Mt*Rv1775 and a putative PhlG homolog of *M. absecssus* (*Ma*PhlG), phloretin hydrolase, which catalyzes C—C bond hydrolysis of a different substrate. Comparison carried out in a similar scheme as in **A.** Superposition of the putative homologs, color-annotated with conserved residues essential for phloroglucol moiety recognition and for phloretin substrate specificity in *Ma*PhlG(47). The structural similarity and conserved Zinc-coordinating and catalytic residues affirm Rv1775 as a *bona fide* C—C hydrolase, potentially with a substrate that includes a phloroglucol moiety, but likely not phloretin. Conservation of structure- and sequence-features characteristic of DinB-like metalloenzymes exemplified by structural homology of Rv2036 and a mycothiol-dependent maleylpyruvate isomerase from *Corynebacterium glutamicum* (*C. glu* MDMPI) **(C-G). (C)** Superposition of Rv2036 structure model and *C. glu* MDMPI (PDB: 2nsf). Conserved Zn^2+^-coordinating (pink) and catalytic residues are highlighted (green). **(D)** Highly conserved residues Arg^222^ (C-terminal domain, Arg^193^ in Rv2036) and Asp^151^ (N-terminal domain, in Asp^124^ Rv2036) are in close proximity (4.22Å), suggesting conservation of their proposed role as inter-domain protein stabilizers(51) **(E)** Spatial conservation of Zn^2+^-coordinating residues of the catalytic triad (Asp and Glu are observed interchangeably) are consistent with conserved catalytic function. **(F)** Surface hydrophobicity of Rv2036 model and 2nsf shows that the hydrophilic core proposed to underlie MDMPI catalysis(51) is relatively conserved. **(G)** Structure-based sequence alignment of Rv2036 and *C. glu* MDMPI with conserved residues were manually annotated according to prior work(51). **(H)** Summary of relevant genomic context potentially informative of function, protein similarity metrics between putative *M. tub* MDMPI homologs and *C. glu* MDPMI, and predicted protein features. All structural images were rendered in PyMol. Structurally homologous sequence alignments are based on TM-align(22) (**<5Å between residues; *<10Å between residues).

**Table 3.**
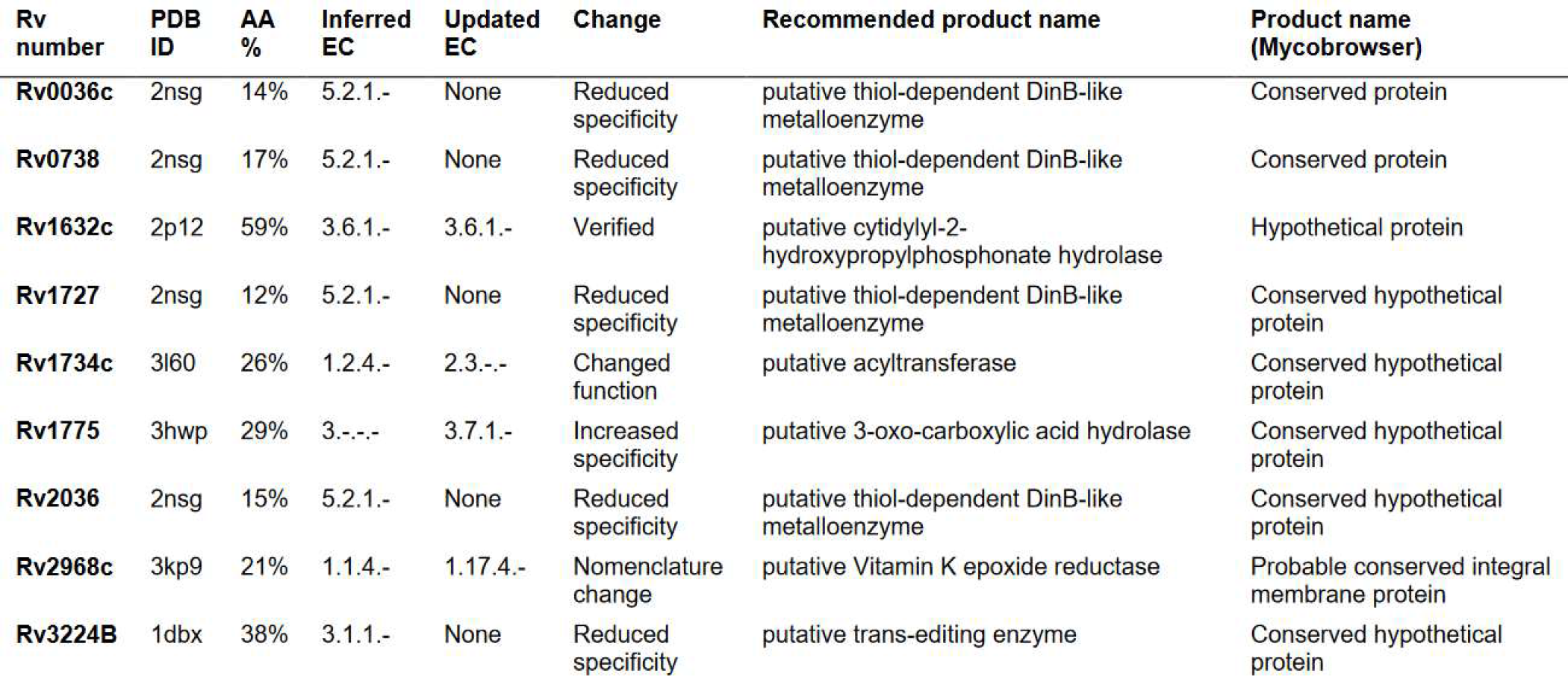
Functional annotation of HHpred-uncorroborated enzymes following manual structural analysis.

In the first example (Fig. 4A-B), manual structural analysis fully corroborates the HHpred-uncorroborated function inferred from structure and extends annotation specificity. Originally, Rv1775 was ascribed putative hydrolase function by our structure-function inference pipeline. Its structural model is globally similar (TM_ADJ_=0.817) to 2,4-diacetylphloroglucinol hydrolase PhlG (EC 3.7.1.24) from *Pseudomonas fluorescens*(46) (PfPhlG) despite only modest sequence similarity (27.6%). Comparison of Rv1775 to PfPhlG (Fig. 4A) and potential mycobacterial homolog (TM_ADJ_=0.753, AA=28.4%) phloretin hydrolase (EC 3.7.1.4) of *Mycobacterium abscessus* (47) (MaPhlG) showed conserved Zn^2+^-coordinating and catalytic residues in the Rv1775 protein model (Fig. 4 A,B). These conserved features suggest Rv1775 encodes a hydrolase acting on C— C bonds (EC 3.7.-.-), an uncommon class of catalytic activity(46). The only subsubclass within EC 3.7.-.-is 3.7.1.-, suggesting Rv1775 is a 3-oxoacid carboxylase.

Although the precise substrate(s) of Rv1775 are indiscernible from structural comparison alone, examining its structure suggests a potential role in lipid metabolism. It shares the phloroglucinol moiety recognition residues conserved across R-phloroglucinol hydrolases but lacks conserved residues required for phloretin hydrolysis (Fig. 4B). This suggests Rv1775 is not a phloretin hydrolase but may act on substrate(s) containing a phloroglucinol moiety or similar aromatic chemical species. Considering reports of *M. tuberculosis* utilizing cholesterol as a carbon source(48), known C—C hydrolytic enzymes in *M. tuberculosis* cholesterol catabolism(49), and gaps in the current understanding of cholesterol catabolism(50), cholesterol ring species are plausible C—C hydrolysis substrates.

In the second example (Fig. 4C-H), we examine one of four HHpred-uncorroborated proteins structurally resembling mycothiol-dependent maleylpyruvate isomerase (MDMPI, a DinB superfamily protein; PDB: 2nsf) of *Corynebacterium glutamicum* (*C. glu* MDMPI). This example illustrates the case when manual inspection corroborates conserved structural features yet precise molecular function remains indiscernible. Manual structural analysis of the putative MDMPI homologs validated that—despite low sequence homology (12-17% similarity)— structural features characteristic of DinB-like enzymes are conserved (shown for Rv2036, Fig. 4). All four putative DinB-like enzyme were highly structurally similar to *C. glu* MDMPI (TM_ADJ_ = 0.63-0.75, Fig. 4C) with a conserved hydrophilic core (Fig. 4F), predicted metal-binding sites (Fig. 4E), retained catalytic triad residues(51) (Fig. 4G), and conserved residues that form a salt bridge between the C- and N-domains(51) of MDMPI (Fig. 4D). However, DinB superfamily proteins comprise several functions (52), making even putative inference of a precise molecular function challenging. Most functionally characterized bacterial DinB-like enzymes are thiol-dependent(52) and the putative MDMPI homologs’ closest structural match was a mycothiol-dependent DinB-like enzyme, suggesting thiol-dependence of these four proteins is probable, likely with mycothiol as the thiol-cofactor (the predominant mycobacterial low molecular weight thiol). We annotated these genes as “putative thiol-dependent DinB-like metalloenzymes” and note as “potential (myco)thiol-dependent S-transferase (EC 2.-.-.-)” (53). For such cases, where structural modelling confidently ascribes protein family and features of structure but not function, integrating knowledge of the function of structural orthologs’, expression data, and genomic context can inform rational speculation about their function (Fig. 4H, *Text S1*).

### Hundreds of annotations inferred by structural similarity

Our structural annotation pipeline inferred function from structure for 400/1,725 under-annotated genes (23.2%, Dataset 1). Structure-derived annotations (mean C-score = 0.39) came from higher quality models (*P* = 1.83x10^-163^, student’s t-test) than proteins without passing matches (mean C-score = -1.91), and more specific annotations tended to come from higher-quality models (*Text S1*, Fig. S5). Structure-based annotation captured putatively shared function for numerous previously unannotated proteins lacking appreciable sequence similarity (Table 4, Dataset 3).

**Table 4.**
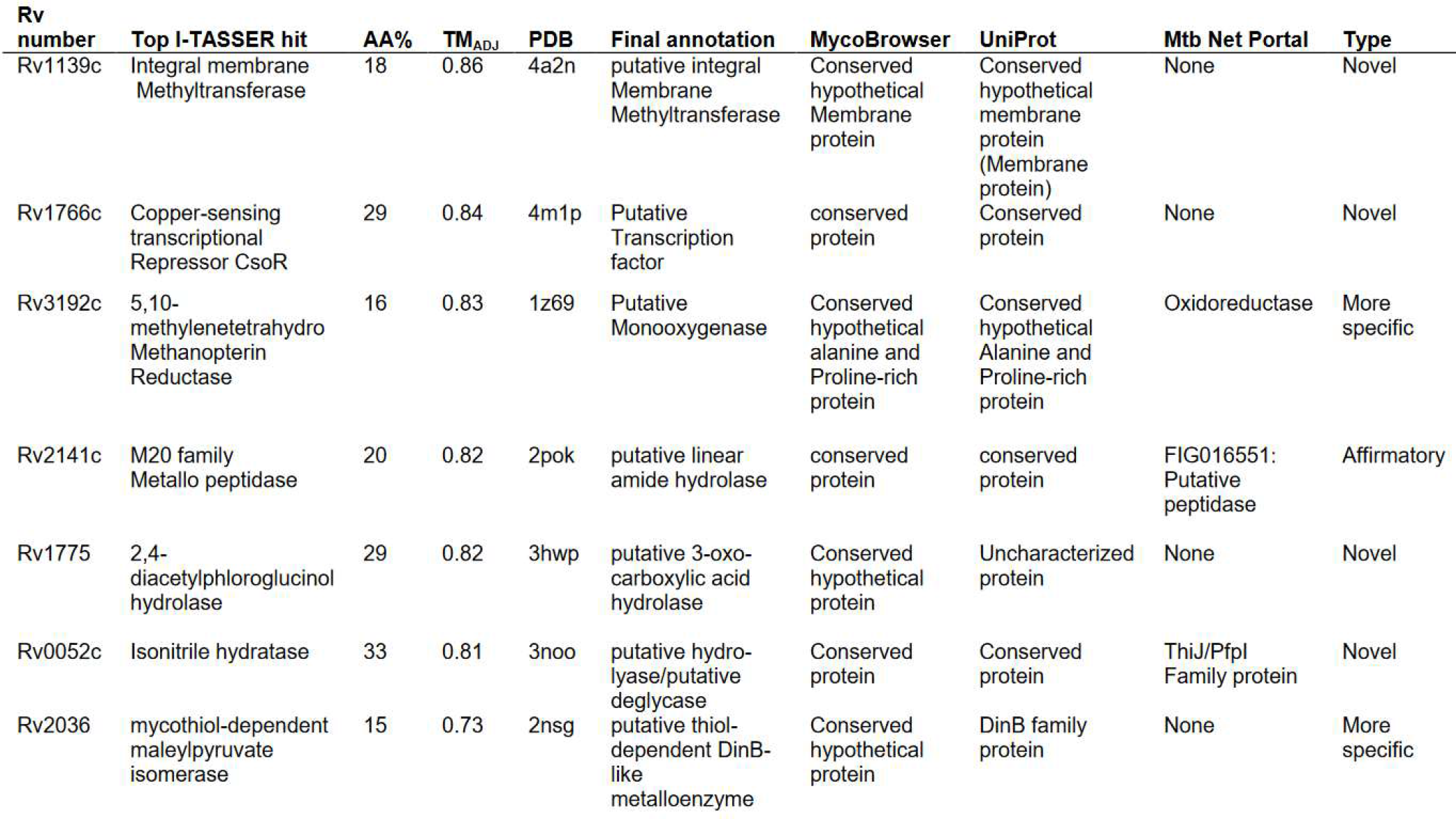
Novel annotations transferred through structural similarity despite low sequence similarity. Selected proteins with modelled structures highly similar to solved PDB structures of known function. Sequence similarities range in the “twilight zone” of sequence similarity, below which remote homology is undetectable by sequence similarity(126). TM_ADJ_ above 0.52 indicates that template and under-annotated gene share structural folds. Annotations from UniProt, Mtb systems Portal, and TubercuList are shown, along with the highest error-adjusted structural similarity match, its identifier (“PDB”), and final product annotation. “Affirmatory” corroborate the annotations in UniProt or Mtb Network Portal. “Novel” are entirely novel annotations to those in UniProt and Mtb Network Portal, while “More specific” are in accord with annotations in other databases but describe product function in greater detail.

These remote homologs and structural analogs include an integral membrane methyltransferase, which can modify of mycolic acids embedded in the *M. tuberculosis* cell wall essential for virulence(54) and redox response-related functions (Rv0052 and Rv3192) critical for enduring host immune defenses in macrophage(55).

### Putative efflux and transport proteins uncovered through structural similarity

Membrane-spanning regions of transport proteins vary in sequence relative to structure far more than globular proteins(12, 56), making them good subjects for structure-based functional inference. Twenty-four proteins were identified as transport proteins and corroborated by HHpred (Dataset 3), including several matches with drug transport proteins (n=8). Eight HHpred-corroborated proteins were not annotated with any transport function in UniProt (Table 5). Rv1510 and Rv3630 exclusively match drug transporters and are uncharacterized across functional databases. Rv3630 mutations have been reported in POA-resistant mutants but no clear causal link was identified(57). Rv1510 is a *Mycobacterium tuberculosis* complex marker in diagnostic assays(58) and its loss-of-function induces autophagy(59), suggesting Rv1510 is an autophagy antagonist important for human-adapted tuberculosis. Verapimil, a potent efflux pump inhibitor, induces autophagy(60), consistent with the putative function of Rv1510 in drug efflux, which could contribute to drug tolerance(58). These putative transporters might contribute to intrinsic efflux-mediated drug resistance and tolerance in *M. tuberculosis* (11). Other putative novel transport proteins may serve important homeostatic roles in the dynamic host microenvironment(61, 62), and could make attractive drug(63) and vaccine(64) targets.

**Table 5:**
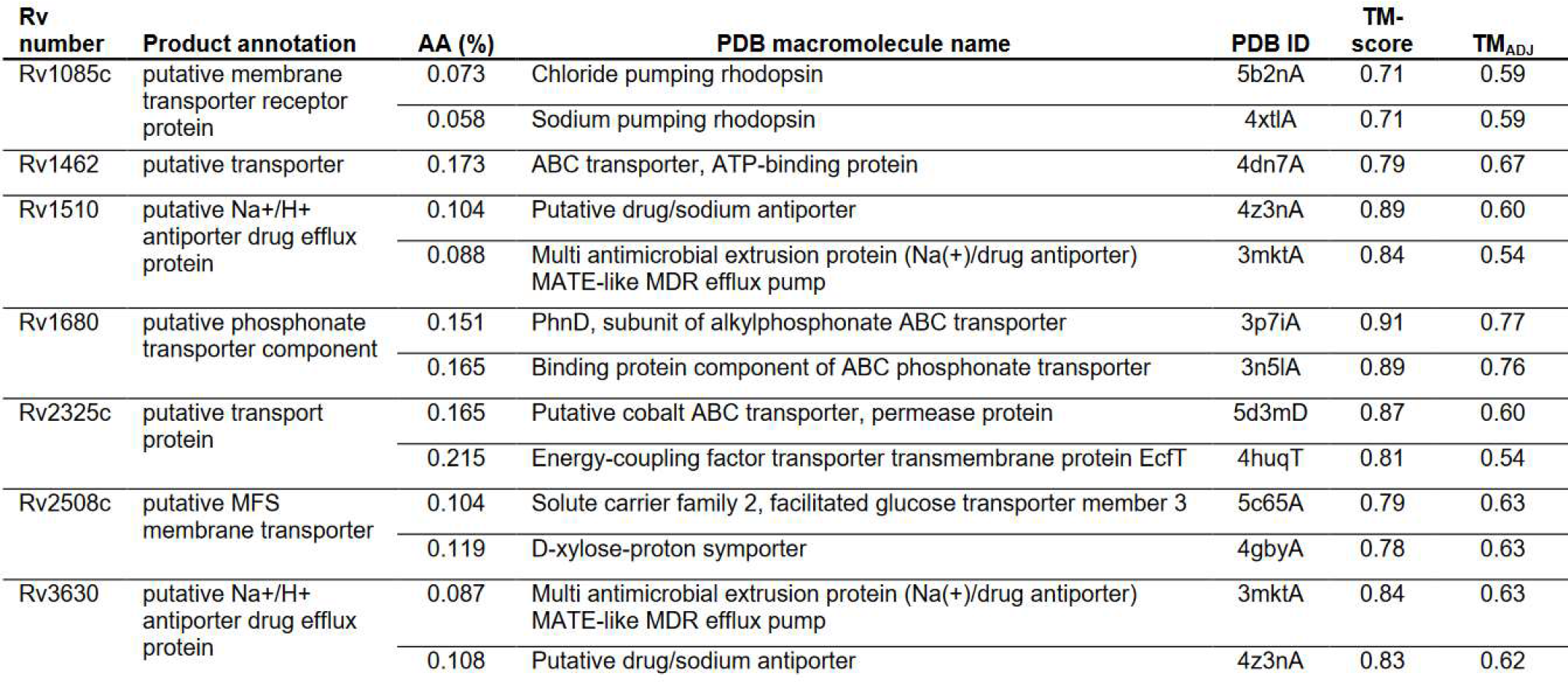
Putative transport proteins. Matches between proteins encoded by under-annotated genes (Locus) and transport proteins structure entries on Protein Data Bank (PDB). Only matches undescribed as transport proteins on Uniprot are included (see Dataset 3E for all such matches). The top two matches are shown, if they exceed adjusted TM-score (TM_ADJ_)) > 0.52 (the TM-score corresponding to matching topologies > 50% of the time). AA% refers to the amino acid identity shared between the aligned region of the protein in *M. tuberculosis* its match on PDB.

| Rv number | Product annotation | AA (%) | PDB macromolecule name | PDB ID | TM-score | TM<sub>ADJ</sub> |
| --- | --- | --- | --- | --- | --- | --- |
| Rv1085c | putative membrane transporter receptor protein | 0.073 | Chloride pumping rhodopsin | 5b2nA | 0.71 | 0.59 |
|  |  | 0.058 | Sodium pumping rhodopsin | 4xtlA | 0.71 | 0.59 |
| Rv1462 | putative transporter | 0.173 | ABC transporter, ATP-binding protein | 4dn7A | 0.79 | 0.67 |
| Rv1510 | putative Na<sup>+</sup>/H<sup>+</sup> antiporter drug efflux protein | 0.104 | Putative drug/sodium antiporter | 4z3nA | 0.89 | 0.60 |
|  |  | 0.088 | Multi antimicrobial extrusion protein (Na<sup>+</sup>)/drug antiporter) MATE-like MDR efflux pump | 3mktA | 0.84 | 0.54 |
| Rv1680 | putative phosphonate transporter component | 0.151 | PhnD, subunit of alkylphosphonate ABC transporter | 3p7iA | 0.91 | 0.77 |
|  |  | 0.165 | Binding protein component of ABC phosphonate transporter | 3n5lA | 0.89 | 0.76 |
| Rv2325c | putative transport protein | 0.165 | Putative cobalt ABC transporter, permease protein | 5d3mD | 0.87 | 0.60 |
|  |  | 0.215 | Energy-coupling factor transporter transmembrane protein EcfT | 4huqT | 0.81 | 0.54 |
| Rv2508c | putative MFS membrane transporter | 0.104 | Solute carrier family 2, facilitated glucose transporter member 3 | 5c65A | 0.79 | 0.63 |
|  |  | 0.119 | D-xylose-proton symporter | 4gbyA | 0.78 | 0.63 |
| Rv3630 | putative Na<sup>+</sup>/H<sup>+</sup> antiporter drug efflux protein | 0.087 | Multi antimicrobial extrusion protein (Na<sup>+</sup>)/drug antiporter) MATE-like MDR efflux pump | 3mktA | 0.84 | 0.63 |
|  |  | 0.108 | Putative drug/sodium antiporter | 4z3nA | 0.83 | 0.62 |

### An updated *M. tuberculosis* reference genome functional annotation

Through manual curation (n=282) and structural inference (n=400), we annotated 623 gene products, reducing under-annotated genes by 36.1%. Including annotated CATH (Class, Architecture, Topology and Homologous superfamily) topologies, functional notes, and Ligand-Binding Sites (LBS) totals 940 (54.5%) with original annotation (Fig. 5B). For genes lacking specific product annotations CATH (Dataset 3L) and LBS assignments (Dataset 3D) can refine functional hypotheses and, in some cases, imply function directly(65). Tetracycline repressor folds (n=17, Dataset 3M), for instance, function nearly exclusively as concentration-dependent transcriptional activators and vary in sequence yet are structurally homogeneous(66). CATH annotations were not used to inform product annotations nor to assign EC numbers in this annotation, however.

**Fig. 5.**
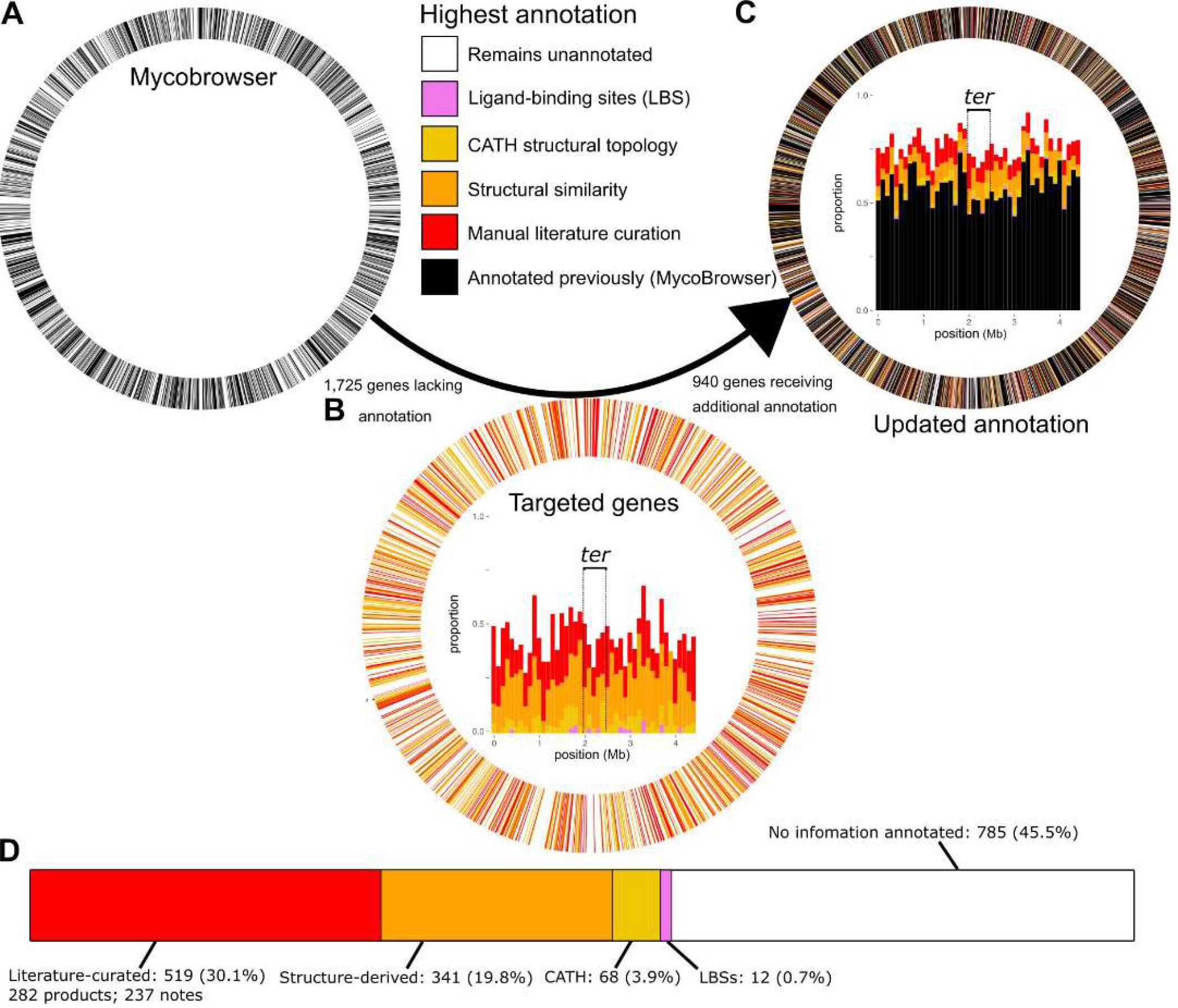
An updated H37Rv functional annotation. (**A-C**) Circos plots illustrating annotation coverage prior to the annotation effort (left) and following it (right), colored according to annotation status. In plots A and C, all 4,031 CDSs are represented as segments of equal width whereas (B) segments the ring into only the 1,725 under-annotated genes. Black genes reflect what was on TubercuList, are considered “annotated”, and are mutually exclusive from the 1,725 under-annotated genes (white). (B) shows only the 1,725 under-annotated genes, whereas A and C include all 4,031 original CDS. Inside the circos rings are stacked bar charts with genes in 100kb bins according to gene start position. The terminal-proximal (± 250 kilobases) region is marked with dashed lines and labelled (*ter*). (**D**) Cumulative number of genes annotated, by annotation type. LBS = Ligand binding site. Percentages refer to under-annotated genes annotated/1,725 initial under-annotated genes. Genes are binned into mutually exclusive categories hierarchically: manually curated product name > structure-derived > literature notes > CATH > LBS. Manually curated and literature notes categories are combined as “Literature-curated” in the visualization. For the purposes of these counts, functional notes from publications implicating many proteins but not clearly establishing function were not counted (e.g. PMIDs 26536359, 21085642, and 22485162).

Our updated annotation provides function for 34.4% (45/131) of genes with hypothetical function identified in a recent systems resource as broadly conserved across *Mycobacteria*(67) (Dataset 3 contains full set). Mycobacterial core genes annotated include functions well-established experimentally, such as essential component of the mycobacterial transcription initiation complex RbpA (https://gitlab.com/LPCDRP/Mtb-H37Rv-annotation/-/blob/master/features/Rv2050.tbl), and others not evident from extant literature but of potential clinical relevance, like the host-directed effector function inferred for Rv3909 (https://gitlab.com/LPCDRP/Mtb-H37Rv-annotation/-/blob/master/features/Rv3909.tbl). These annotations came in similar numbers from published experimental evidence (n=21) and structural inferences (n=24).

Updated annotations distribute across all segments of the chromosome (Fig. 5A) and implicate efflux proteins (Table 5), metabolic functions (Fig. 6), virulence factors, and functions key to survival during infection (Table 6) and under drug pressure (Table 7). Yet many under-annotated genes remain without products or functional notes assigned (n=785). Of these 785 remaining under-annotated genes (Dataset 1), 190 have quality models (C-score > -1.5) but lacked annotations meeting inclusion criteria. Meanwhile, 182 of those remaining have product annotations in UniProt or *Mtb* Network portal. Remaining still, however, are 466 under-annotated genes with neither quality structure nor functional annotation in these databases. These genes frequently cluster consecutively along the genome (99 genes across 23 clusters, Dataset 1G), forming syntenic blocks of unknown function. Genomic context suggests several of these clusters have roles in virulence and drug tolerance (Dataset 1G).

**Fig. 6.**
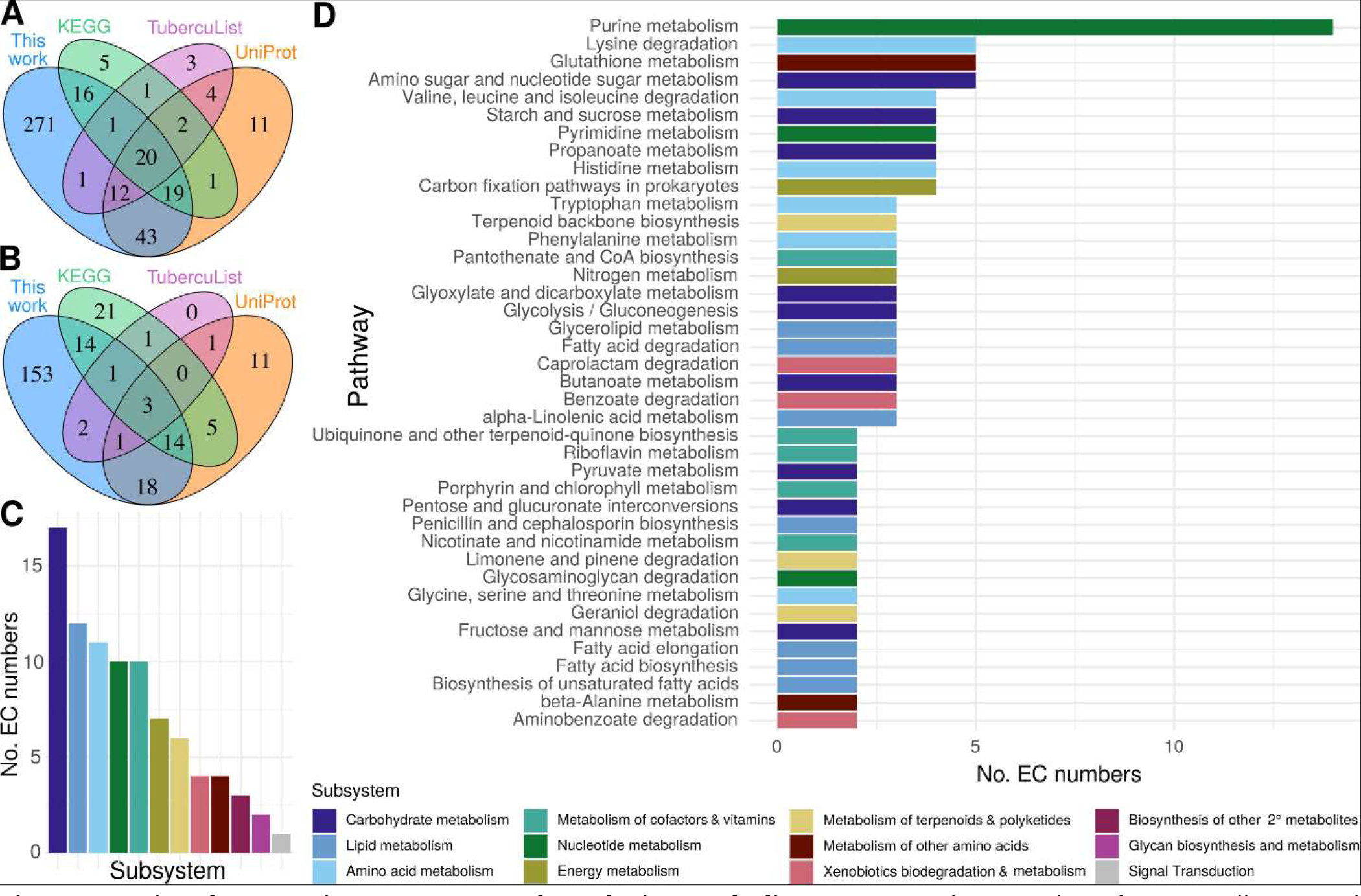
Functional annotations across *M. tuberculosis* metabolism. Annotated EC numbers for manually curated and structure-inferred products were compared with annotations for each under-annotated gene in popular databases (**A**) Set analysis of under-annotated genes (UAG) with an EC number assigned in this study compared to popular databases. (**B**) Novelty of EC numbers for UAG annotated in this study with respect to popular databases. (**C & D**) Distribution of EC numbers annotated across KEGG (**C**) Subsystems and (**D**) Pathways. Generic KEGG subsystems are depicted. All pathways with at least three genes have the number of EC numbers displayed. For subsystems with no pathways with three or more genes, the highest total pathway is displayed.

**Table 6.** Updated annotations add functional knowledge to genes required for optimal fitness during TB infection. Source annotation is the annotation listed by Bellerose and colleagues(83) and new annotation derived from the current project. Protein data Bank identifiers (PDB ID) of the protein structures matching H37Rv protein models are listed for structure-based annotations. PubMed IDs are listed for the papers from which functional annotations were manually curated.

| Rv number | Source annotation | New annotation | PubMed ID | PDB ID |
| --- | --- | --- | --- | --- |
| *Literature annotations* |  |  |  |  |
| Rv1205 | hypothetical protein | riboside monophosphate phosphoribohydrolase | 25728768 |  |
| Rv2018 | hypothetical protein | probable antitoxin VapB/Antigen | 28066388,<br>23140854 |  |
| Rv2272 | transmembrane protein | Probable gamma delta T-cell activator | 23389928 |  |
| Rv2525c | tat pathway signal sequence | probable peptidoglycan hydrolase | 16952959,<br>25869294,<br>25260828 |  |
| Rv2923c | hypothetical protein | probable Osmotically induced bacterial protein C (OsmC, a probable peroxide reductase) | 22088319 |  |
| Rv3632 | membrane protein | Putative flippase | 21030587 |  |
| Rv3763 | lipoprotein LpqH | adhesin/antigen LpqH | 16098710,<br>12594264 |  |
| Rv3788 | hypothetical protein | probable secondary channel binding-factor of RNA polymerase | 22194445 |  |
| *Literature & Structural inference* |  |  |  |  |
| Rv0191 | integral membrane protein | Putative efflux pump | 25690361,<br>12520088,<br>22132058 | 5c65, 4gby |
| Rv1433 | exported protein | Probable I,d-transpeptidase LdtMt3 | 24041897 | 3tur, 3vae,<br>4jmn |
| Rv1769 | hypothetical protein | T-cell antigen/putative aldehyde-lyase | 26853625,<br>15102765 | 4v15 |
| Rv3722c | hypothetical protein | probable serine hydrolase, probable aspartate transaminase |  | 3ppl |
| *Structural inference* |  |  |  |  |
| Rv0047c | hypothetical protein | Putative transcriptional regulator |  | 1yg2a, 3l9f |
| Rv0259c | hypothetical protein | Putative lyase |  | 2jh3, 4ccs |
| Rv0323c | hypothetical protein | Putative hydro-lyase/Putative linear amide hydrolase |  | 1q7t, 5cgz |
| Rv0449c | hypothetical protein | Putative oxidoreductase |  | 2ive, 1sez,<br>3nks, 3i6d, 3lov |
| Rv0767c | hypothetical protein | putative transcription factor |  | 3mnl |
| Rv1085c | hemolysin-like protein | putative membrane transporter receptor protein |  | 5b2nA, 4xtlA |
| Rv2052c | hypothetical protein | putative endodeoxyribonuclease |  | 3igh |
| Rv2160A | hypothetical protein | putative transcription factor |  | 2hyj |
| Rv3226c | hypothetical protein | putative peptidase |  | 2icu |
| Rv3433c | hypothetical protein | putative hydro-lyase/putative isomerase |  | 2ax3 |
| Rv3719 | hypothetical protein | putative amide-bond oxidoreductase |  | 3dq0, 2exr,<br>4o95 |

**Table 7.**
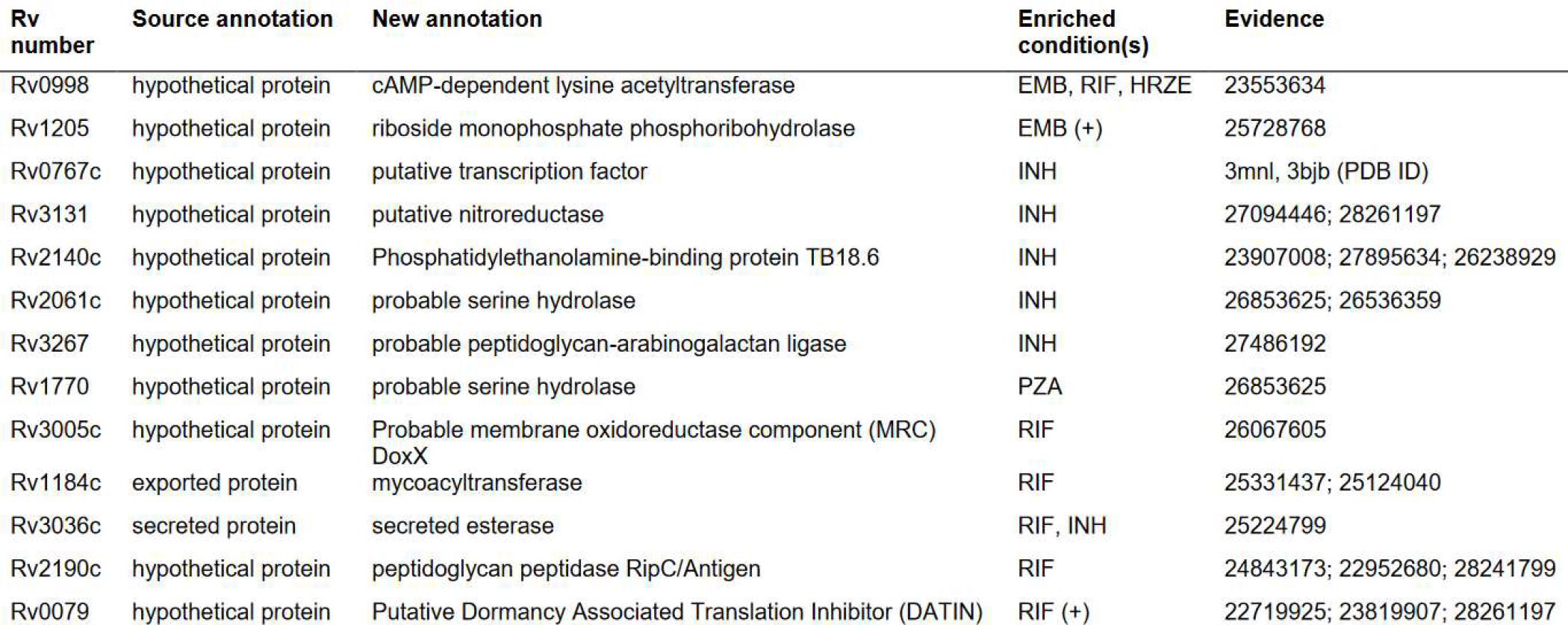
Updated annotation enriches functional interpretation of under-annotated genes affecting drug sensitivity. Source annotation is the annotation listed by Bellerose and colleagues(83). Enriched conditions are the drugs’ exposure under which differential mutant abundance was observed. Sources of updated annotation are listed in the evidence column. INH=Isoniazid; RIF=Rifampicin; EMB=Ethambutol; PZA=Pyrazinamide; HRZE=Combination regimen of INH, RIF, PZA, and EMB. “(+)” indicates enrichment observed at multiple time points.

Genes remaining without any form of annotation (Fig. 5D) were overrepresented (*P* = 0.0011, odds ratio=1.35, Fisher’s Exact) near the terminus (±250 kilobases from half the genome length) of the chromosome (*ter*-proximal genes, Dataset 1H). An even stronger bias for uncharacterized genes can be seen for genes transcribed opposite the direction of replication (*P* = 1.14 x 10^-7^, odds ratio=1.53; Fisher’s exact). To ensure that circumstantial factors such as PE/PPE or insertion element density were not accounting for the apparent orientation and spatial trends across the chromosome, we removed all PE/PPE and insertion seq and phage genes and repeated the analysis. The trend strengthened for the *ter*-proximal gene bias (*P* = 0.0034, odds ratio=1.44; Fisher’s exact) and decreased only marginally (*P* = 2.53 x 10^-6^, odds ratio=1.49; Fisher’s exact) for the orientation bias.

These biases are consistent with three previously noted trends that could influence the likelihood of gene characterization. First is the general trend of decreased gene expression as a function of distance from the *oriC* in bacteria(68). On average, highly expressed genes are more amenable to functional characterization. Second is the strong bias for symmetric inversions around the terminus(69), particularly in Actinobacteria(70). Hypotheses leading to experimentally determined functions are often informed by orthology, which can be inferred by conserved synteny between species(71). Common inversions around the terminus can disrupt this synteny with increased frequency. Disruption can occur globally—through moving across the chromosome by inversion—and locally, by inversion boundaries interrupting operons or other syntenic features. Third, genes transcribed opposite the direction of replication frequently collide with the replication machinery, making them more mutable than genes with transcription and replication co-oriented(72). This increases the likelihood of weakened promoters or loss-of-function mutations evolving *in vitro* for genes non-essential in H37Rv. One potential confound is that genes encoding virulence/toxin proteins are enriched on the lagging strand(72). As these genes operate in the context of infection, they are challenging to functionally characterize, which may contribute to the observed enrichment of uncharacterized genes on the lagging strand.

Turning our attention to metabolism, 381 under-annotated genes were annotated with EC numbers (Dataset 1I Methods & Materials), over two-thirds of which were absent from other databases (Fig. 6A,B). Fully specific (4^th^ EC digit) EC numbers (n=92) were ascribed to 85 genes. These newly annotated reactions span diverse metabolic pathways and subsystems (Fig. 6, Dataset 1J), many implicated in mediators of *M. tuberculosis’* virulence such as lipid and polyketide and terpene metabolism(73-75), which are integral to the unique composition of the mycomembrane. Proteins of these pathways have important immunity-subverting functions(76) at the host-pathogen interface(77). For instance, terpenes play an immunomodulatory role early in *M. tuberculosis* infection and phagosomal maturation(78-80), are potential agonists of antibiotics for TB treatment(81), and include cell membrane surface-expressed molecular species unique to *M. tuberculosis*(82). The numerous carbohydrate-metabolizing products (Fig. 6D) may identify alternative metabolic pathways in *M. tuberculosis* and aid in gap-filling efforts in *M. tuberculosis* metabolic reconstructions.

### Integration with recently published functional screens

Next, we assessed how much novel functional information our annotation added to ambiguously or hypothetically annotated genes from a recent transposon mutagenesis study that sought to identify specific bacterial functions limiting drug efficacy during a mouse model of infection(83). We assessed two sets of genes identified in the study. In the first set of under-annotated genes—those newly reported to as essential for optimal growth in mouse infection—one-third (23/69) could be updated by our annotations (Table 6). Fifteen were structural inferences, demonstrating the value structure-based inference of putative function where the difficulty of recapitulating complexities of the host environment challenges functional elucidation through experiment. Notably, following its inference based on structure, Rv3722c has since been confirmed to indeed encode an aspartate transaminase(84) and Rv1085c has been found likely not to encode hemolysin(85), substantiating the structure-derived functional annotations in Table 6.

Our annotation functionally described 13/27 under-annotated genes affecting drug sensitivity (Table 7). Notably, some genes affecting drug sensitivity have published functions consistent with the mechanism of action of the drug of interest but listed without annotation. For instance, the authors noted cell wall permeability as a central theme among genes dictating sensitivity to rifampicin (RIF), and disruption of Rv2190c—a peptidoglycan hydroase—rendered mutants hypersusceptible to RIF, consistent with an effect on cell wall permeability. Others (e.g. Rv1184c) were unannotated in their primary data, but their functional ties was discussed in the text, suggesting the function was curated from literature. Our updated annotation centralizes such functional knowledge.

### Structural models enable functional interpretation of novel PZA-resistant mutants

Next, we applied our annotations prospectively to a new resistance screen, querying the molecular basis of pyrazinamide (PZA) resistance in *M. tuberculosis*. PZA is a cornerstone of modern tuberculosis therapy, yet the mechanism by which it exerts its anti-tubercular activity remains elusive. PZA is a pro-drug that must be converted to its active form pyrazinoic acid (POA) by a mycobacterial amidase(86). While multiple explanations for POA action have been proposed(87–89), many of these models have not held up to scrutiny(90-92). Recently, several groups have shown that POA either directly or indirectly disrupts mycobacterial coenzyme A (CoA) biosynthesis(93-95),. Identification of novel resistance mechanisms could shed additional light on the elusive action of this drug. Thus, a library of 10^5^ transposon-mutagenized *M. tuberculosis* H37Rv was used to select for POA resistant isolates. While the frequency of spontaneous resistance to POA is approximately 10^-6^, the frequency of resistance from our transposon mutagenized library was 10^-3^. Four mutant strains chosen for further characterization of drug resistance profile and transposon insertion site (Fig. 7) showed insertions in genes of unknown function. Each of these strains showed ≥2-fold resistance to PZA and POA (Fig. 7A-D) and no change in INH susceptibility (Fig. 7A-D) when compared to wild-type H37Rv.

**Fig. 7.**
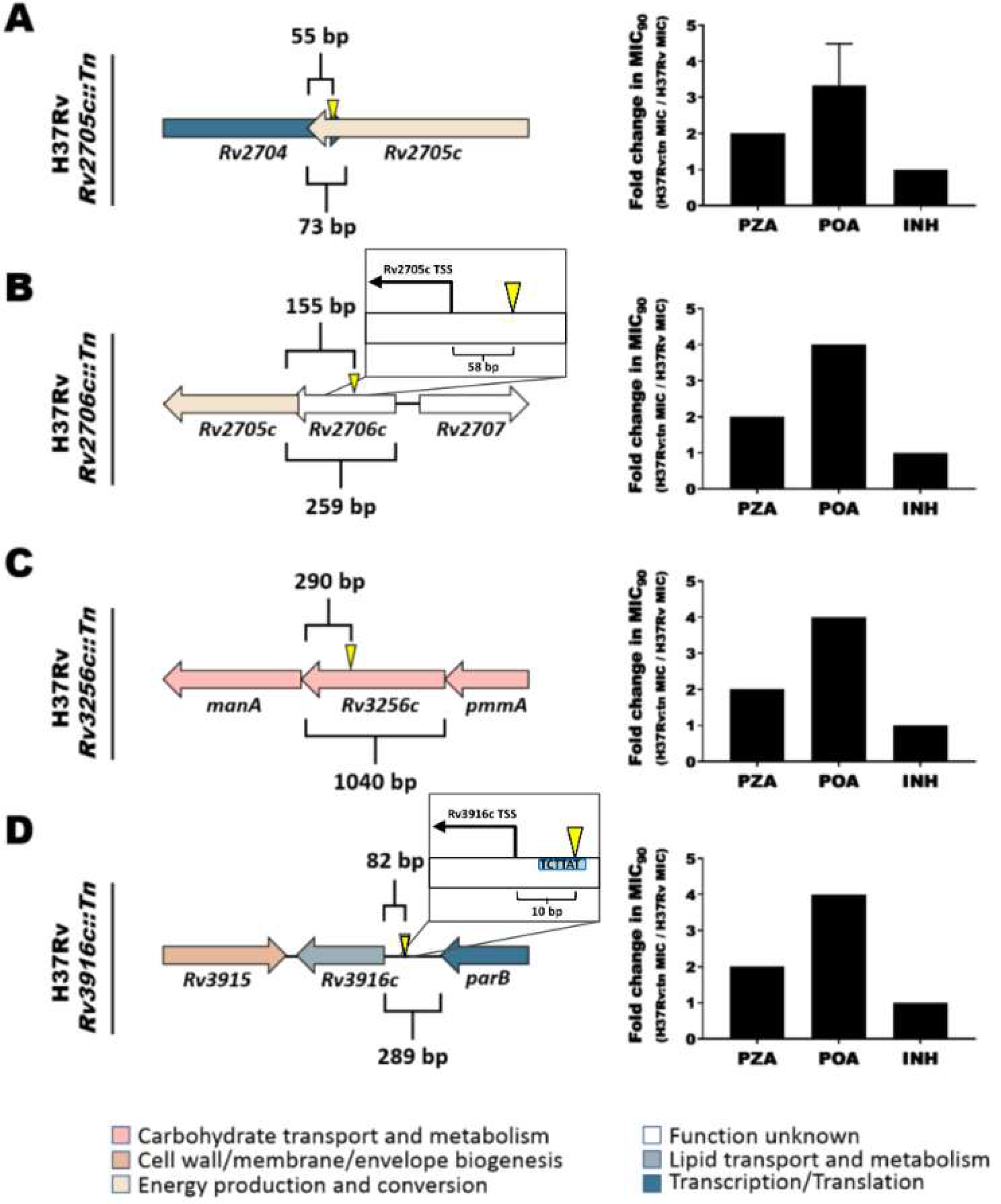
Annotation of genes involved in pyrazinamide resistance. A library of *M. tuberculosis* H37Rv transposon insertion mutants was used to select for strains that were resistant to POA. The transposon insertion sites were mapped and the strains were characterized for their susceptibility to PZA, POA, and INH in comparison with wild-type H37Rv (MIC_90_: 50 µg/mL PZA, 200 µg/mL POA, 0.0625 µg/mL INH). (A) H37Rv *Rv2705c::Tn*. (B) H37Rv *Rv2706c::Tn*. (C) H37Rv *Rv3256c::Tn*. (D) H37Rv *Rv3916c::Tn*. Error bar depicts standard deviation across triplicates. Pop-outs in (B) and (D) depict the transposon insertion sites relative to experimentally determined Transcription Start Sites(127, 128) (TSS). The insertion in (D) interrupts a TANNNT Pribnow Box (blue), destroying the Rv3916c promoter. Error bars are shown when there was a deviation in the calculated MIC90 across triplicates.

To interpret how the interrupted genes might contribute to PZA resistance, we inspected the structural and functional data available from our I-TASSER results (Figs. 8 & 9). PZA is a structural analog of nicotinamide(96), suggesting the putative nicotinamide binding domain of Rv2705c (Fig. 7A) may interact directly with PZA or POA. While it remains difficult to confidently annotate Rv2706c (Fig. 7B), considering its position immediately upstream of Rv2705c it may alter PZA sensitivity by influencing expression of Rv2705c.

**Fig. 8.**
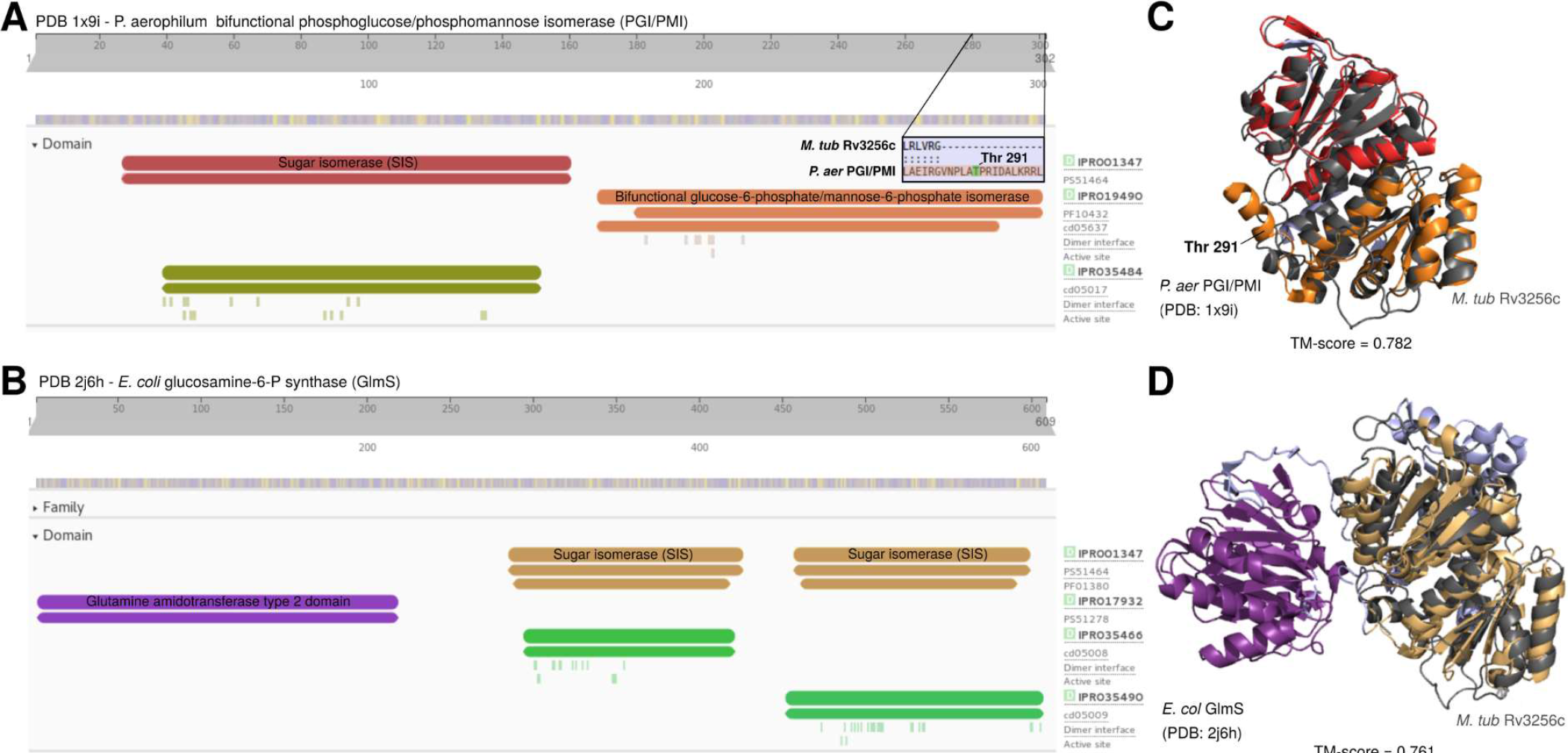
Structural analysis informs specific functional hypotheses for the basis of PZA resistance in the *Rv3256c::Tn* mutant. (**A-D**) Structural analysis identifies Rv3256c as a sugar isomerase (SIS) domain-containing protein likely involved in phosphosugar metabolism or its regulation. InterPro functional domains are displayed for the two strongest structural matches of Rv3256c, (**A**) *P. aerophilum* bifunctional phosphoglucose/phosphomannose isomerase (*P. aer* PGI/PMI) and (**B**) *E. coli* glucosamine-6-P synthase (*E. col* GlmS). The InterPro domains labelled in A and B are mapped onto the 3D structures of (**C**) *P. aer* PGI/PMI and (**D**) *E. col* GlmS. Rv3256c (charcoal) modeled protein structure is optimally superposed to each of its matches. Rv3256c is structurally homologous to the SIS domains of *E. col* GlmS and *P. aer* PGI/PMI and exhibits the alpha-beta-alpha sandwich fold of SIS(129). The pop-out in A (**<5Å between residues) and labelled residue in C show the threonine residue essential for isomerase activity in *P. aer* PGI/PMI (Thr^291^) and other PMI homologs. The Thr^291^-containing region appears to be absent from Rv3256c. Likewise, Rv3256 lacks a glutamine amidotransferase domain homologous to *E. col* GlmS. From this structural evidence, we conclude that Rv3256c is a SIS domain protein putatively involved in phosphosugar metabolism and/or its regulation. Structural images were rendered in PyMol. Structurally homologous sequence alignments from TM-align(22).

**Fig. 9.**
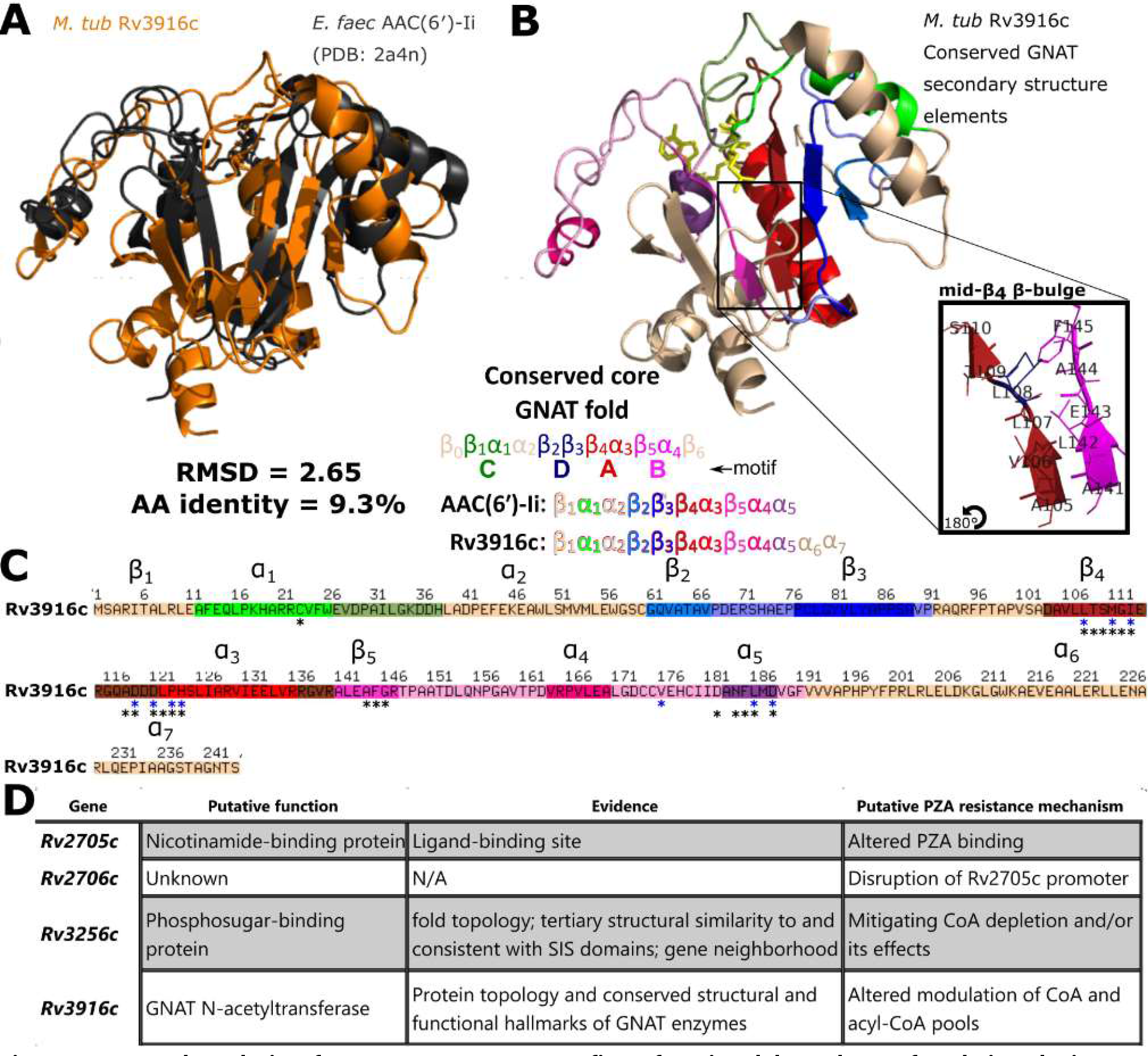
Structural analysis of transposon mutants refines functional hypotheses for their role in PZA resistance. (**A-C**) Structural analysis supports Rv3916c as a General control non-repressible 5 (GCN5)-related N-acetyltransferases (GNAT). Rv3916c matches exclusively comprised GNAT enzymes with low sequence similarity, typical among homologous GNAT enzymes (130). (**A**) Rv3916c (orange) superposed with its closest structural match (PDB 2a4n, charcoal), *Enterococcus faecium* aminoglycoside 6′-N-acetyltransferase (*E. faec* AAC(6′)-Ii). (**B**) *M. tuberculosis* Rv3916c structural model (top) and secondary structure topology (bottom), colored according to the four core conserved GNAT folds (wheat = poorly conserved secondary structure element) and secondarily (tones of the primary colors) by secondary structure elements in *E. faec* AAC(6′)-Ii (as defined by (131)) with which Rv3916c structurally aligns. All secondary structure elements of *E. faec* AAC(6′)-Ii are present in Rv3916c, with two gratuitous alpha-helices in its C-terminal arm (which is not well-conserved among GNAT enzymes(130)). The distinctive β-bulge (pop-out) within the β4 strand (red, bulge residues colored blue) characteristic of GNAT enzymes is present in Rv3916c, diverting β4 away from β5 to create the chasm where the acetyl-coenzyme A (yellow) is predicted to bind (**C**) Primary sequence of Rv3916c colored according to the scheme described in (B). Asterisks mark predicted acetyl-CoA binding residues (black) and residues structurally aligning to known CoA-interacting residues (blue). All known CoA-interacting residues from *E. faec* AAC(6′)-Ii are conserved in Rv3916c and all predicted acetyl-CoA binding sites coincide or directly flank demonstrated sites of CoA interaction. The presence of the features in Rv3916c suggest it is a GNAT enzyme. GNAT enzymes catalyze transfer of an acyl moiety from an acyl-CoA to various substrates (130) making Rv3916c a probable acyl-CoA acetyltransferase and implicating _P_Rv3916c::Tn in CoA pool modulation model of PZA resistance. (**D**) Summary of functional hypotheses for PZA resistance-conferring transposon mutants. All structural images were rendered in PyMol. Structurally homologous sequence alignments are based on TM-align(22).

Rv3256c structurally resembles multiple phosphosugar isomerases—particularly phosphoglucose (PGI) and phosphomannose (PMI)—and Glutamine-fructose-6-phosphate transaminases (GlmS). Rv3256c has neither the conserved residues essential for PMI/PGI catalysis (97) (Fig. 8A,C) nor the glutamine amidotransferase domain required for GlmS activity (98) (Fig. 8B,D), effectively ruling out these functions. The common structural feature among these functionally disparate matches is a sugar isomerase (SIS) domain (Fig. 8). SIS domain is a phosphosugar-binding module(99), implicating Rv3256c in phosphosugar metabolism or its regulation. Rv3256c lacks the HTH domain common to RpiR-like SIS domain proteins(99) that regulate phosphosugar metabolism genes, refuting the possibility of a RpiR-like transcriptional regulatory function. Flanking Rv3256c (Fig 7C), however, are mannose donor biosynthesis genes—Rv3255c (a PMI) and Rv3257c (a phosphomannomutase). In *M smegmatis*, Rv3256c overexpression decreased cell-surface mannosylation(100), consistent with a role in regulating phosphosugar metabolism, though a specific molecular function for Rv3256c remains unclear. Presumably, the role played by Rv3256c in phosphosugar metabolism is disrupted in the Rv3256c::Tn mutant. This disruption may alter composition of acyl-CoA/CoA pools (e.g., through disrupting/promoting acylation of cell wall constituents) or the metabolic and cell wall restructuring response under CoA depletion(93) induced by PZA treatment.

Rv3916c structurally resembles numerous Gcn5-related N-acetyltransferase (GNAT) proteins and exhibits structural, topological, and local features characteristic of GNAT enzymes (Fig. 9). Perhaps most tellingly, Rv3916c has a predicted acetyl-CoA binding site consistent with known GNAT enzymes (Fig. 9B,C). While the acyl donor and substrate of Rv3916 remain unclear, the conservation of these structural features involved acyl-CoA interaction in functionally characterized GNAT superfamily proteins strongly suggest that Rv3916 is a GNAT N-acetyltransferase. This putative function implicates the PRv3916c::Tn mutant in the CoA-depletion model of POA action. Destruction of the Rv3916c promoter would reduce its expression, in turn altering acyl-CoA pool modulation. Structure-function insights gleaned from these structural models inform specific functional hypotheses for these mutants’ role in PZA resistance and demonstrate how the provided structural data can enrich the interpretation of large-scale screens and generate specific functional hypotheses.

## Discussion

Functional genome annotation is critical for interpreting the deluge of omics data generated by emerging high throughput technologies. Here, we devised procedures to systematically curate annotations from published literature and infer putative function through structure-based function inference and applied them to annotate the *M. tuberculosis* virulent type strain and primary reference genome, H37Rv. We curated annotations for hundreds of proteins with published functions lacking from common resources (Table 2), a quarter of which were absent from all five annotation resources examined, highlighting the importance of community-specific manual curation. To complement these manually curated annotations, we built a structural modeling and functional inference pipeline (Figs. 1-4), calibrated it to include confident annotations (Fig. 1), orthogonally validated it with established remote homology detection methods (Fig. 3). Through structure-function inference we annotated hundreds of genes (Fig. 5), including dozens of potential transport proteins, resistance genes, and virulence factors (Dataset 1). Elucidating the determinants of *M. tuberculosis’* survival under drug pressure and within the context of infection are chief objectives of tuberculosis research. Integrating this updated functional annotation with new (Figs. 7-9) and published (Tables 5 & 6) functional screens showed it can aid understanding the genetic basis of *M. tuberculosis’* resistance to drug pressure and infection-like conditions. The structural models provided a rational basis for functional hypotheses of the molecular basis of resistance for four novel PZA-resistant strains with mutations in otherwise unannotated genes (Figs. 7-9). In particular, the Rv3256c and Rv3916c mutants implicate CoA homeostasis in PZA resistance (Figs. 8 & 9), linking them to the CoA depletion model of PZA mode of action described recently for other PZA-R mutants (93). The functional interpretation of TnSeq mutants afforded by this resource informs hypotheses for the mechanistic basis of these mutants’ PZA resistance for investigation in future work.

Our systematic approach to manual literature curation has limitations. First is the time and attention from researchers with specialized knowledge required for manual literature curation. To mitigate this limitation in the future, contributions from the TB research community can be submitted, and will be incorporated with standardized criteria and structured ontologies(24, 28). A second limitation is the inevitable subjectivity of the curator. We addressed this by requiring two curators review each paper independently and provided explicit guidelines for what evidence warrants annotation with the degree of confidence connoted systematically by qualifying adjectives.

Other limitations arise from the scope of our annotation. First, we only curated functions for 1,725 of over 4,000 ORFs in *M. tuberculosis*. Products that did not meet our criteria for inclusion may have useful functional characterizations excluded by our approach. Second, we only searched for the locus tag during curation. While most publications include locus tag, some do not and therefore some experimental characterizations may remain unannotated. Last, we searched literature back from 2010, as TubercuList updated continuously through March 2013, and we assumed annotations to that point were captured. However, the absence of dozens of characterizations from all resources suggests some findings prior to 2010 may remain unintegrated. Despite these limitations, the numerous genes we curated that were absent from all frequented annotation sources are now centralized in an single updated annotation that is clear in source and confidence level, in a consistent and extendable format.

Multiple factors contribute to error and bias in resolving protein structure and function. These factors fall unevenly across protein classes and families(101), making them challenging to account for. Considering this while designing our structure-based inference pipeline, we favored simple, interpretable inclusion criteria, coupled with downstream quality assurance measures. Future work more focused on customizing inclusion criteria optimized for features of protein structure or function may improve prediction accuracy. Our simplified approach let us circumvent accounting for these biases explicitly, which would require further methods development and introduce additional bias if not executed carefully.

Our structural approach to functional inference also has limitations. First, it depends on the input sequence. We took amino acid sequences as provided by TubercuList without accounting for the impact of known, uncorrected sequencing errors(102) or corrections to amino acid sequences proposed by UniProt curators. Furthermore, some genes have multiple translation initiation sites, or isoforms(103), but we considered one sequence per gene. Second, our approach compares global, rather than local, structural similarity and can be challenged by functionally diverse folds(104), and proteins with dynamic active sites(105) or context-specific conformation and activity(106). Our empirically-driven inclusion criteria (Fig. 1) and quality control measures helped to mitigate false positive annotations (Figs. 3 and 4). In future analysis of structural models, emerging methods that capture functional conservation distributed across primary and tertiary structure may identify functionally informative protein features missed by our approach. Promising approaches include direct coupling analysis(107), statistical coupling analysis(108), Bayesian partitioning with pattern selection(109), and structurally interacting pattern residues’ inferred significance(110). Third, proteins from model organisms and humans are overrepresented among crystallized structures on PDB (https://www.rcsb.org/) (111). This adds bias toward inferring function from these proteins. Finally, the structure-based annotations should be interpreted as tentative, since inclusion criteria required similarity implying >50% (“putative”) or >75% (“probable”) likelihood of being correct. Structure-based annotations should be viewed accordingly, as well-informed hypotheses rather than established truth.

Over half of structure models (871/1,711) were low-quality (C-score < -1.5)(21) (Dataset 1). Several phenomena may challenge effective modeling of these under-annotated genes: 1) No proteins of similar folds have been solved; 2) The protein is highly disordered(112); 3) These are multi-domain proteins that need to be split into individual domains(14); 4) Sequencing errors; 5) gene coordinate misannotation(102); 6) Pseudogenization. We suspected reason 3 as a major factor, considering we did not attempt to break up multi-domain polypeptides into their constituent domains(14). However, the protein length distributions of proteins of high (greater than -1.5) and low (below -1.5) C-scores were similar (*Text S1*, Fig. S5C), which suggests the presence of multiple domains was not a primary cause of poor models. Each of the other reasons likely contributes to some extent, but reasons 1 and 2 are most troublesome for PE/PPE genes and other protein classes specific to *Mycobacteria*.

I-TASSER failed to produce models for 14 under-annotated genes (Dataset 1). Six of these sequences are pseudogenes and the remaining 8 belong to PPE or PE_PGRS gene families, which are especially prone to sequencing errors and intrinsically hypervariable(113). Although we ascribed putative functions for some PE/PPE genes the function of most remains unclear. Far fewer PE/PPE proteins (20/166, 12.0%) than non-PE/PPE genes (518/1,559, 33.2%) met inclusion criteria for structure-based annotation (*P* = 2.13x10^-9^, odds ratio = 0.276; Fisher’s exact). This likely owes partly to their intrinsic disorder and partly to their specificity to the *M. tuberculosis* complex(114), which limits the number of homologous structures in PDB with known function, challenging accurate structural modeling and structure-based functional annotation. Moreover, PE/PPE and other effector proteins require precise metabolic contexts or immunological cues, precluding observation of their function *in vitro*. Characterizing function for these genes will require high-throughput biochemical assays and development of techniques that directly assay or precisely reconstruct host microenvironments; formidable challenges, indeed. In the meantime, carefully designed and caveated inferential methods can make valuable surrogates and streamline candidate prioritization for experimental confirmation or more comprehensive *in silico* analysis.

Systematically curated literature and structure-derived annotations are available at https://gitlab.com/LPCDRP/Mtb-H37Rv-annotation. Researchers can file issues to report future published characterizations and submit merge requests to incorporate future functional characterizations. These methods can continue to furnish annotations as functional characterizations are published in the primary literature, structure-function relationships in PDB expand, *M. tuberculosis* gene product functions are determined, and sequence-structure-function prediction tools become more resource-efficient.

## Author contributions

SJM, AE, and FV conceptualized the overarching research goals and aims; SJM, AE, DG, AMZ, NK, and CKC developed the manual curation protocol; SJM, DG, AMZ, NK, and CKC manually curated annotations from literature, and independently validated annotations; SJM, AE, DG, AMZ, NK, and CR developed and tested code for data processing and analysis; AE and SJM developed the structural comparison workflow; SJM, AE, and ND developed and executed the structural QC workflow; NAD and ADB conceptualized the PZA resistance experiments; NAD isolated and characterized the PZA resistant mutants; SJM, AE, DG, AMZ, NAD, and ND prepared the manuscript; SJM, DG, AMZ, and NAD prepared figures and tables; SJM and FV managed the project.

## Materials and Methods

Additional detail on methods are provided in *Text S1*

### Data availability

We provide final annotations in common machine (GFF3) and human (Dataset 1) readable formats, including EC numbers, GO terms, CATH topologies, and product name annotations. Annotations in the GFF3 are defined by our inclusion criteria. PDB templates with structures similar yet below our criteria are provided in Dataset 3 (top 3 PDB templates for each under-annotated gene) and Dataset 3 (all matches where TM_ADJ_ > 0.52 (Equation 2, *Text S1*) and/or µ_geom_ > EC3 (putative) threshold). I-TASSER results and model protein structures for 1,711 under-annotated genes are freely accessible at https://tuberculosis.sdsu.edu/structures/H37Rv/ including functional predictions by COFACTOR, predicted ligand binding sites, local secondary structure confidence (B-factor), and other quality and similarity metrics.

### Manual curation protocol

All publications mentioning each of the 1,725 under-annotated gene were independently evaluated for annotation-worthy functional characterization by two researchers and quality checked by a third for format and protocol compliance (*Text S1*, Fig. S1). Qualifying adjectives were defined by evidence quality and systematically assigned to connote annotation confidence. Notes relevant to function but insufficient to assign product name were also annotated (*Text S1*, Fig. S1).

Precision benchmarking. We designed procedures and inclusion criteria to maximize precision (Equation 1) and minimize “Overannotation”(101): Only annotations with 50% or greater precision were included, regardless of source. Whereas other metrics had applicable precision benchmarks (*Text S1*), EC number and GO terms did not. We assessed how precision of EC number and GO term predictions (Equation 2) correlated with similarity metrics. We evaluated which I-TASSER metrics were most predictive of precision (Equation 1) through logistic regression (*Text S1*, Fig. S2).

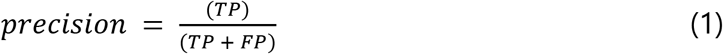

where T*P* = “True Positive”, and F*P* = “False Positive.

We gauged sequence similarity by amino acid identity (AA%) and structural similarity by Template Modeling Score (TM-score). TM-score describes structural similarity from 0 and 1. It represents the average root-mean squared deviation across all atoms in the predicted structure with respect to the PDB template model, normalized to remove apparent deviation arising falsely due to local differences(14, 115) (Methods & Materials), allowing proteins of different length to be compared(115).

To base inclusion criteria off precision (Equation 1), we regressed against sequence and structural similarity metrics: amino acid identity, C-score, TM-score (Equation 1, as calculated by Zhang and Skolnick(115)), and the geometric mean of TM-score and AA% (µ_geom_) against precision of EC number assignment in a positive control set of 363 *M. tuberculosis* genes with known function but unknown structure. Because EC numbers and GO terms encode the same fundamental information, although GO terms have many false negatives and were relatively underpowered (*Text S1*, Fig. S2), we included both according to the same criteria: µ_geom_ values corresponding to 50% (“putative”) and 75% (“probable”) precision for each tier of specificity (Fig. 1).

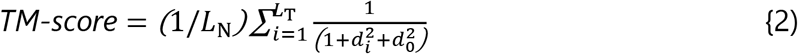

where *L_N_* is protein length, *L_T_* is the length of the aligned residues to the template, *d_i_* is the distance of the *i*^th^ pair of residues between two structures after an optimal superposition, and 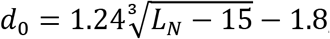, as described by Zhang *et al.*, normalizes for protein length(21). TM-score measures the difference between *predicted* structure and known structure of the putative homolog/analog on PDB. Since we are interested in the similarity between the *true* (unknown) structure and its putative homolog/analog on PDB, we used an adjusted TM-score, “TM_ADJ_”. TM_ADJ_ subtracts from TM-score the expected difference in TM-score between the modeled protein structure and its (unknown) true structure (See Equations 2 and 3 and accompanying text in *Text S1* for additional discussion of this rationale).

### Training data selection

We combined 200 randomly selected *M. tuberculosis* protein sequences with known function with 163 manually annotated under-annotated genes with “probable” or higher annotation confidence (Dataset 1) to form a set of training genes. We extracted EC numbers and GO terms that were marked as experimentally verified in UniProt(116) from the 363 training genes.

### Annotation inclusion criteria

Structure-inferred annotations comprised product names, GO terms, EC numbers, CATH topologies, and LBS. We systematically transferred EC numbers and GO terms according to µ_geom_ thresholds corresponding to 50% and 75% precision (Fig. 1). LBS and CATH predictions were included according to previous precision benchmarks(21, 117). CATH annotations were retrieved using the REST API of PDB for structure matches surpassing the TM-score corresponding to 50% precision, after correcting TM-score for expected modeling error (S2 Text). Under-annotated genes with quality models (C-score > -1.5) and a TM-score greater than 0.85 and/or µ_geom_ meeting the inclusion criteria for putative EC 3rd digit (0.374, corresponding to a precision > 0.5), and further criteria based on aligned portions and method of UniProt annotation of the PDB template (*Text S1*). Because transport proteins are more conserved in structure than in sequence relative to globular proteins(12) we weighted structural similarity more heavily than AA% in their inclusion criteria: Transport protein annotations were transferred if (i) greater than 90% of the PDB structure implicated in transport aligned with the under-annotated gene model, and (ii) structural similarity exceeded the threshold for CATH topology transfer. All analyses were implemented in R(118).

### Product naming protocol

To translate transferred GO terms and EC numbers into product names we converted GO terms that describe enzymatic activity into EC numbers by searching EXPaSY ENZYME. Product names were converted from EC numbers (including those derived from GO terms) using the ENZYME.dat file from the EXPaSY database (*Text S1*, Figs. S3 and S4). GO terms that mapped to multiple EC numbers, were merged at the most specific level at which they converged (e.g., 3.2.1.5 and 3.2.2.4 would resolve to 3.2.-.-). When GO terms did not map to an EC numbers, we translated sufficiently descriptive GO terms into product names (e.g., “DNA binding transcription factor activity” is sufficiently descriptive whereas “pathogenesis” is not). Product names for PDB matches lacking GO or EC annotations were determined manually. Transport proteins were named with lower specificity than their PDB matches (e.g. “transport protein” instead of “Na+/H+ antiporter”), unless (i) all three strongest PDB matches converged on a more specific description and (ii) TM-score exceeded 0.85 for at least one of the three, the name in common between the three strongest matches was transferred. LBS predictions and the residues predicted to coordinate binding (Dataset 3D) can be interpreted as being at least 60% likely to be true(23), though most have greater confidence.

For proteins annotated only with structure-based functional inferences that had EC number annotation modified by HHpred-filtering, or had multiple EC numbers corroborated by HHpred, the implicated structural homologs were inspected manually, and spurious or infeasible annotations were pruned. Reasons for pruning EC numbers include cases where one of the implied catalytic functions was exceedingly unlikely (such as eukaryotic proteins with bacterial homologs that had evolved distinct, non-overlapping functions) or there was a clear reason for a false positive (such structural alignment with a multifunctional protein to only one of the functional domains). Additional annotation specificity was added in rare cases, where HHpred results strongly corroborated evidence from structural alignment that alone did not meet inclusion criteria for specific annotation. Rv3433c exemplifies such cases. It was annotated with EC 4.2.1.- and EC 5.-.-.- and both were corroborated by HHpred. Upon inspection, the annotators noted that the top hits from structural alignment and HHpred were a mixture of EC 5.1.99.6 and EC 4.2.1.136 proteins and bifunctional proteins encoding both catalytic functions. The portions aligning to the respective EC functions were mutually exclusive, and Rv3433c was of similar length to characterized bifunctional enzymes including both functions. In this case, EC numbers were updated to full specificity and the product name was changed from “putative hydro-lyase/putative isomerase” to “putative Bifunctional NAD(P)H-hydrate repair enzyme”

### Comparison with other databases

To assess the novelty of manual product annotations, we compared our annotation for each under-annotated gene with the corresponding entry on UniProt(116), Mtb Network Portal(9) (which included annotations from TBDB(5)), PATRIC(6), RefSeq(36), BioCyc(119), and KEGG(120). Comparisons were performed programmatically where possible, and systematically otherwise.

### Enzyme commission number assignment

We assigned EC numbers to under-annotated genes with experimentally verified enzymatic activity using that assigned by the source article’s author when compliant with IUBMB standards. Otherwise, we manually assigned one using the official IUBMB database(25).

HHpred filtering of structure-inferred functional annotations. All proteins with functions assigned solely by structural inference were run through HHpred, searching against the PDB70 and ECOD databases, limiting maximum number of hits to 1000, and using default parameters for the searches. All HHpred results were filtered, and only hits where Prob > 0.95 were retained for downstream analysis. Each function assigned to a protein was evaluated separately (e.g. a bifunctional protein could have one function culled and the second function retained). Annotations were evaluated differently depending on whether they had a corresponding EC number or now. All annotations with an EC number assigned were evaluated programmatically and retained to the degree of EC specificity matched by HHpred hit(s). Annotations without corresponding EC numbers were evaluated manually, independently, by two curators. Each curator screened all HHpred hits and evaluated whether HHpred hits supported the assigned function wholly, entirely, or not at all. In cases where function was partially supported, each curator submitted a suggested product name change. After evaluating all proteins, the curators reconciled any disparate assignments. Functions entirely uncorroborated by HHpred that passed the Ramachandran plot analysis filtering step were subsequently evaluated to determine whether the structural similarity used to infer function had substantial evidence warranting a transferred annotation. Original structural inferences were either discarded, retained, or modified at the discretion of the curators. For an annotation to be accepted, curators verified that model proteins were not threaded on low-complexity proteins, checking whether regions underlying function of the structurally solved protein structurally aligned to the protein model being annotated, and for conservation of any known residues or structural motifs essential for function.

### Ramachandran plot analysis

To evaluate structure model quality we computed the fraction of the residues in “most favored” regions, “additionally allowed” regions, “generously allowed” regions and “disallowed regions”, via the PROCHECK server (121). The .pdb file containing the atomic coordinates of each model protein structure of interest was uploaded to PROCHECK and proportions of residues in each regional favorability classification were extracted from the “results summary” file and collated into tabular format for further analysis. To determine a threshold for including structures not corroborated by HHpred on the basis of quality protein structure, we assessed the distributions of “most favored” region residues between proteins wholly uncorroborated by HHpred and those fully corroborated by HHpred with maximally specific EC numbers (Fig. 3).

### Structural model visualization and annotation

Protein structure models (.pdb files) from I-TASSER and solved protein structures form The Protein Data Bank were visualized and annotated with pymol (https://pymol.org/).

### Comparison with annotations from recent functional screens

Tables S2 and S4 from a recently published Transposon mutant functional screen(83) were downloaded and the intersection of their locus tags and the under-annotated gene set of this study went on for further analysis. Genes with annotated functions by Bellerose and colleagues(83) were filtered out. PE/PPE genes and Transcriptional regulatory proteins were also excluded, as the novelty comparison was determined according to product name, which typically remains generic for these two classes of proteins even upon updated functional information.

### Bacterial strains and growth media

*M. tuberculosis strain* H37Rv was a gift from W.R. Jacobs, Jr. of the Albert Einstein College of Medicine. Strains were grown in Middlebrook 7H9 medium (Difco) supplemented with 10% (vol/vol) oleic acid-albumin-dextrose-catalase (OADC) (Difco), 0.2% (vol/vol) glycerol, and 0.05% (vol/vol) tyloxapol.

Characterization of POA resistant strains. Strain H37Rv was mutagenized with the *mariner*-based transposon(122, 123). Approximately 10^5^ independent transposon mutagenized bacilli were plated on Middlebrook 7H10 medium supplemented with 10% (vol/vol) OADC (Difco), and 0.2% (vol/vol) glycerol with 50 μg/mL of pyrazinoic acid (POA) (Sigma). Resistant mutants were selected from the plates containing 50 μg/mL POA. The initial isolates were plated on 7H10 medium supplemented with 10% (vol/vol) OADC (Difco), and 0.2% (vol/vol) glycerol either containing 400 μg/mL POA or no drug after their initial isolation to confirm their POA resistance prior to the more detailed drug susceptibility testing. (123, 124) Transposon insertion sites were identified as previously described.

The antimicrobial drug susceptibility was determined by assessing the minimum concentration of drug that was required to inhibit 90% of growth (MIC_90_) relative to a no drug control. Growth was assessed by measuring OD_600_ of cultures after 14 days of incubation at 37°C. Drug susceptibility testing for PZA and POA was carried out in 7H9 broth supplemented with OADC, glycerol, and tyloxapol (pH 5.8) as indicated above. INH MIC_90_ determinations were carried out in medium with the same composition at pH 6.8.

## ACKNOWLEDGEMENTS

We extend our gratitude to Dr. Vikram Alva for running a large set of proteins through HHpred in batch mode on our behalf. We thank Sarah Radecke, Derek Conkle-Gutierrez, and Matthew Onorato for their extensive review of the manuscript and helpful comments. We would also like to acknowledge Dr. Yusuke Minato for his contributions to PZA experiments. This work was funded by grants from National Institute of Allergy and Infectious Diseases (NIAID Grant No. R01AI105185 to FV and R01AI123146 to ADB). SJM, AMZ, DG, AE, NK, CR, ND, CKC and FV were supported by R01AI105185. SJM was also supported by scholarships from a National Science Foundation DUE Training Grant to FV (0966391). ADB and NAD were funded by R01AI123146. NAD was also supported by NHLBI (HL007741). The funding bodies had no role in the design of the study or in collection, analysis, and interpretation of data or in writing the manuscript.

## Supplementary Figures

**Fig. S1.**
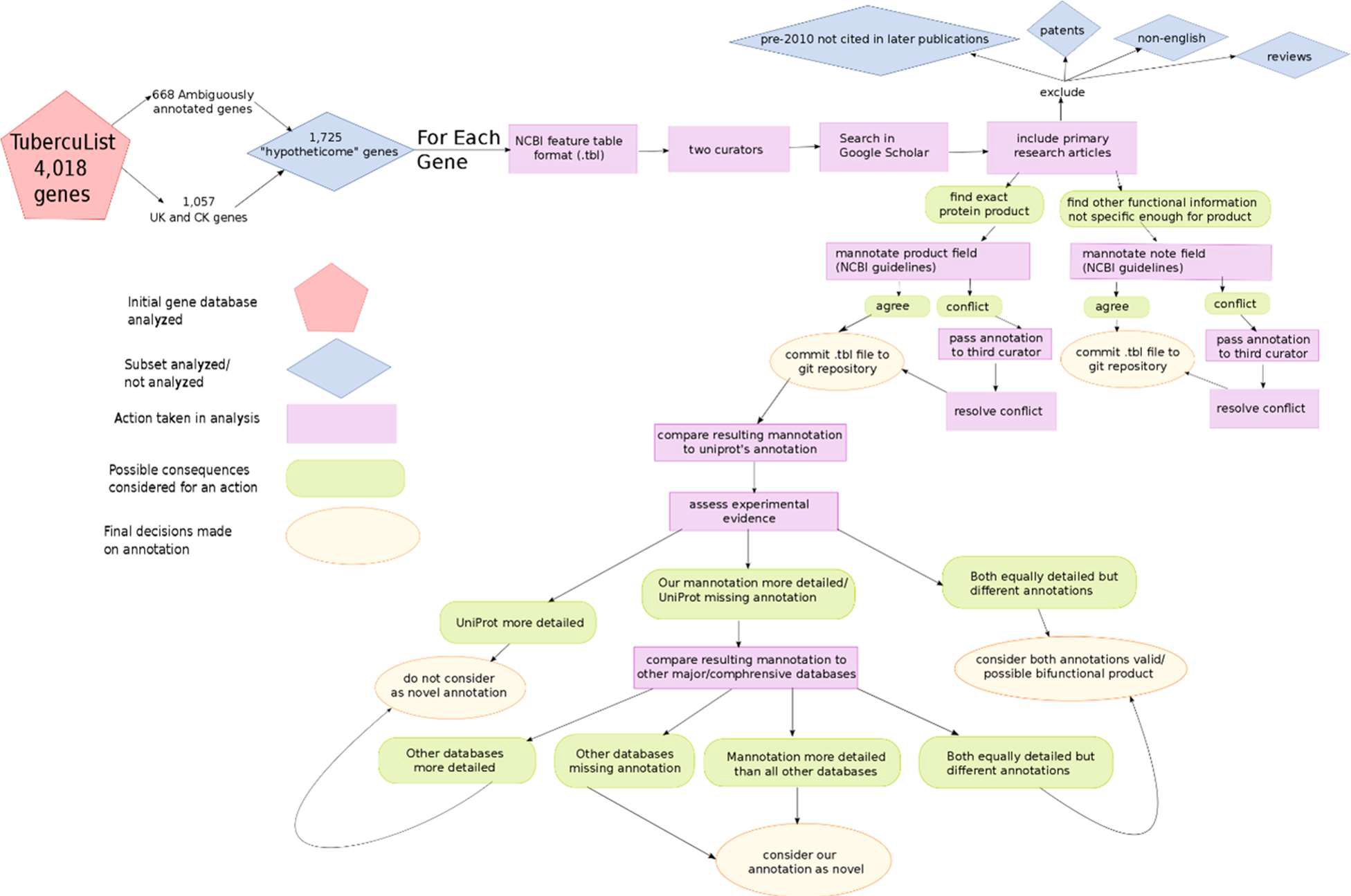
Information flow for producing annotations from literature curation. An initial extraction of the existing annotation every “conserved hypothetical” and “unknown” protein from TubercuList totaled 1,057 unannotated protein-coding genes. Additionally, a 668 “ambiguous set” was manually determined from the annotations on TubercuList, and these annotations were extracted and combined with the 1,057 hypothetical and unknown proteins to give a total of 1,725 under-annotated genes. These genes were then searched in Google Scholar, and pertinent articles were analyzed for annotation information, which was recorded in NCBI’s Table File Format (.tbl extension) for each gene, one file per gene. Every gene annotated with a novel product was compared to annotation in other databases (Results). Decisions were made based on the criteria depicted above.

**Fig. S2.**
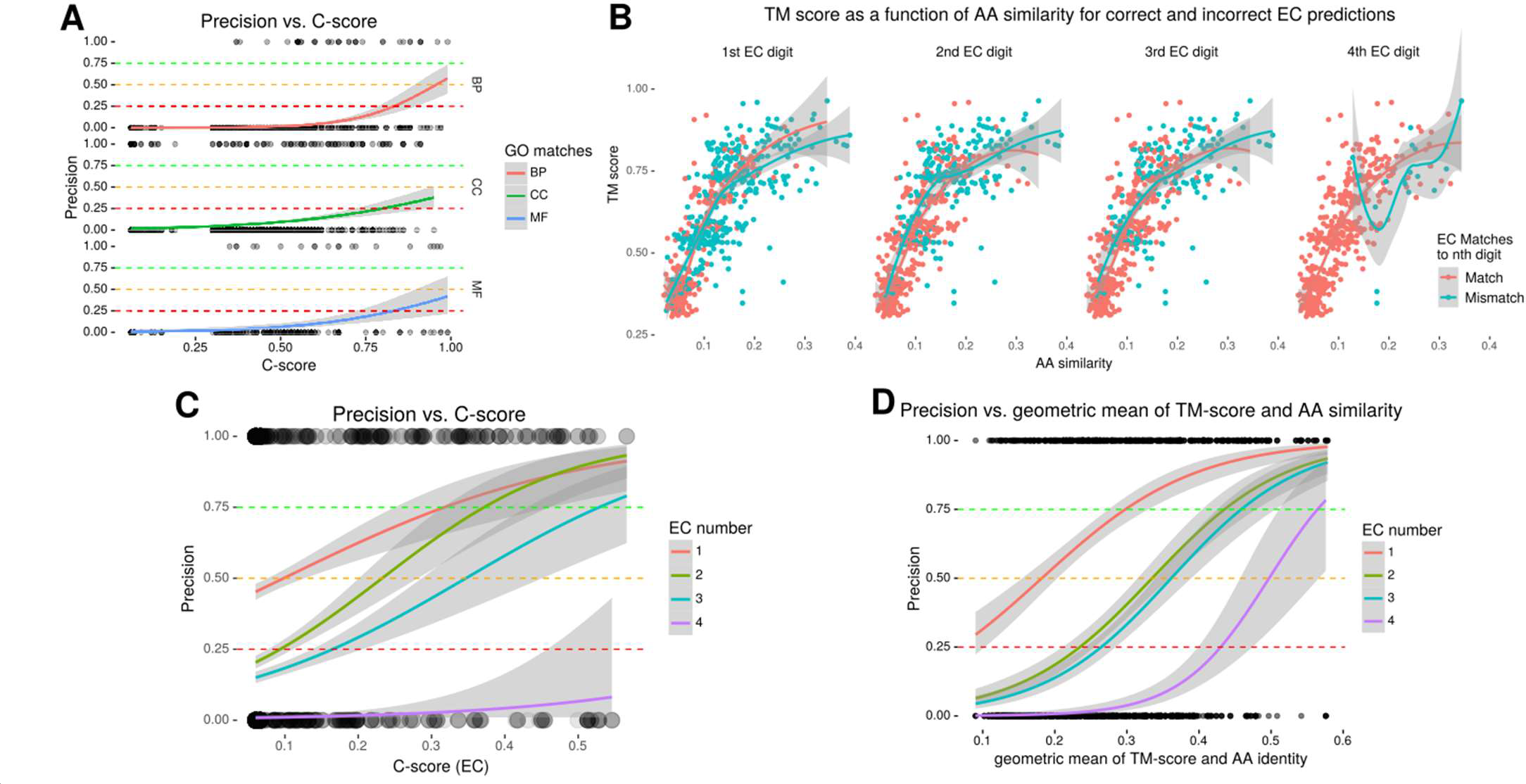
Relating precision to similarity metrics. A-D display binomial regressions on similarity metrics. **(A)** Precision of GO term prediction as a function of C-scores from COFACTOR. Precision of GO term predictions are plotted as a function of C-score for each of the three ontologies (Biological process, BP; Cellular Component, CC; Molecular Function). The circles at the bottom and top of each plot are individual data points (representing 0 and 1 for incorrect and correct, respectively at a particular value). The circles are rendered at 10% opacity to visually depict observation density. For benchmarking, only templates with C-scores above -1.5 were included, as structural predictions with lower confidence are unlikely to reflect correct protein topology(21). **(B)** TM-score and amino acid sequence identity (AA%) colored by correctness in the sample data. Dots’ position indicates structural similarity (TM-score, y-axis) and AA% (x-axis) between modeled structure and PDB entry, and their color indicates concordant (red) or discordant (blue) EC number, to the specificity indicated by the pane label. **(C)** Function of EC number precision using C-scores from the structure-to-function platform COFACTOR(23). Horizontal lines indicate the cutoffs used to set the thresholds for hierarchical incorporation. **(D)** Similar to **A & C**, but using the geometric mean of Amino acid identity and TM-score. This is the same data plotted in Figure 1 of the main text, and the model we selected for determining inclusion criteria.

**Fig. S3.**
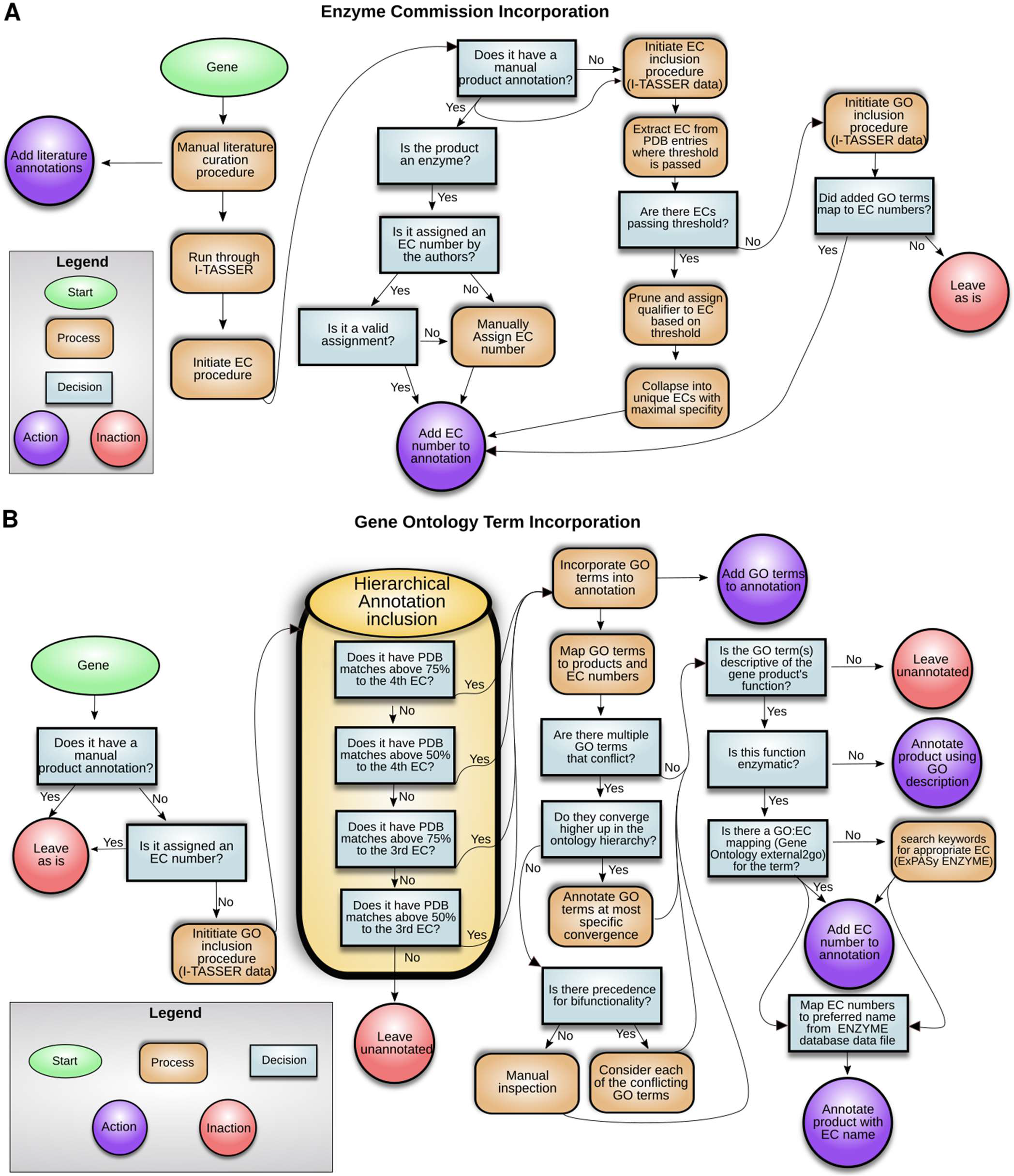
Structure-based annotation inclusion protocols. **A) Enzyme Commission (EC) number inclusion protocol.** The flow of processes and decisions each protein model was subjected to for determining EC based annotations from structural homologs. Most processes and decisions were implemented in a fully automated manner, but some corner cases had to be resolved manually. These manual cases were handled algorithmically, or by previously established procedures where possible. For example, when EC numbers had to be assigned manually, the procedures put forth by IUBMB were consulted and followed directly(25). **B) Gene Ontology (GO) term inclusion protocol.** The flow of processes and decisions each modeled protein structure was subjected to for determining GO based annotations from structural homologs/analogs. All processes and decisions were implemented in a fully automated manner, up until product assignment, those of which did not map to EC number had to be resolved manually.

**Fig. S4.**
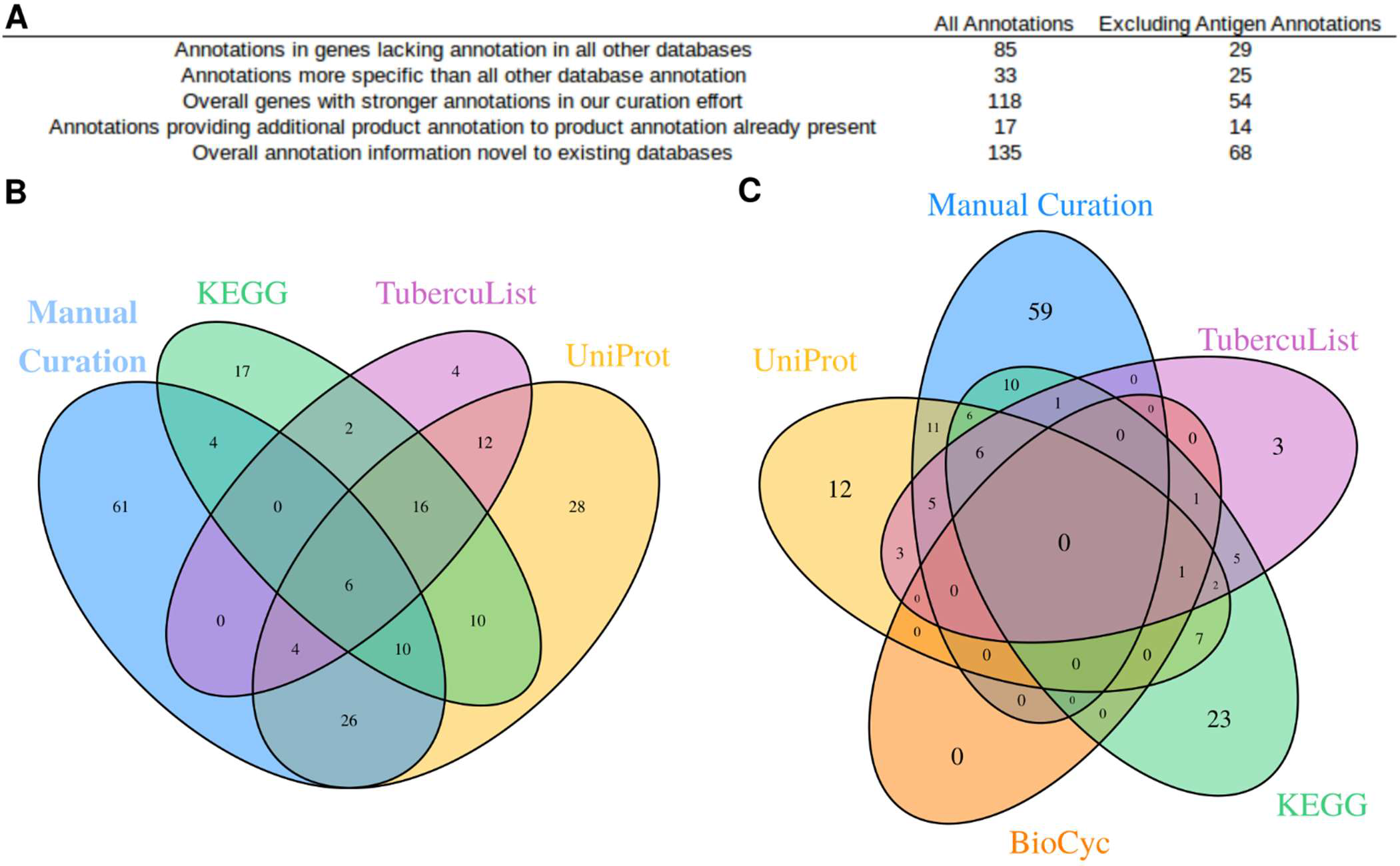
Comparison of annotations to existing databases. (**A**) To assess the novelty of manual product annotations, we compared each to their counterparts on UniProt(116), Mtb Network Portal(9) (which included annotations from TBDB(132)), PATRIC(6), RefSeq(36), BioCyc(119), and KEGG(120). We obtained the UniProt, Mtb Network Portal, and RefSeq annotations for each of our genes with new product annotation programmatically. For each under-annotated gene, we determined whether the database annotations agreed or disagreed with our manually curated annotations. For annotations that agreed, we recorded which source had the more descriptive annotation. Annotation from our literature curation absent in the other databases were considered novel gene annotations. If the annotations disagreed, we considered our annotation a candidate for additional gene product annotation, since both our annotation and those in other databases may describe true functions (bifunctional/moonlighting proteins). Existence of functional annotations for these genes were tallied for each database to assess their comprehensiveness and identify discrepancies between them. Furthermore, genes unannotated in any of the listed databases, but with annotations assigned in this study, were identified and enumerated. EC number assignments were also compared among the databases (Results). Of these databases, BioCyc and UniProt are the most comprehensive for GO term annotations, while UniProt and Mtb Network Portal have the fewest hypothetical proteins (Table 1). **(B-C) EC number annotation in the manual curation effort compared to widely used databases.** Manually curated annotations were compared with those in the databases in Table 1 to identify the presence or absence of EC number in GUF annotations. (**B**) Set of under-annotated genes annotated with an EC number in each of the five databases compared. The non-overlapping segments indicate the number of under-annotated genes annotated uniquely in that database. (**C**) Set of unique EC numbers across under-annotated for each database.

**Fig. S5.**
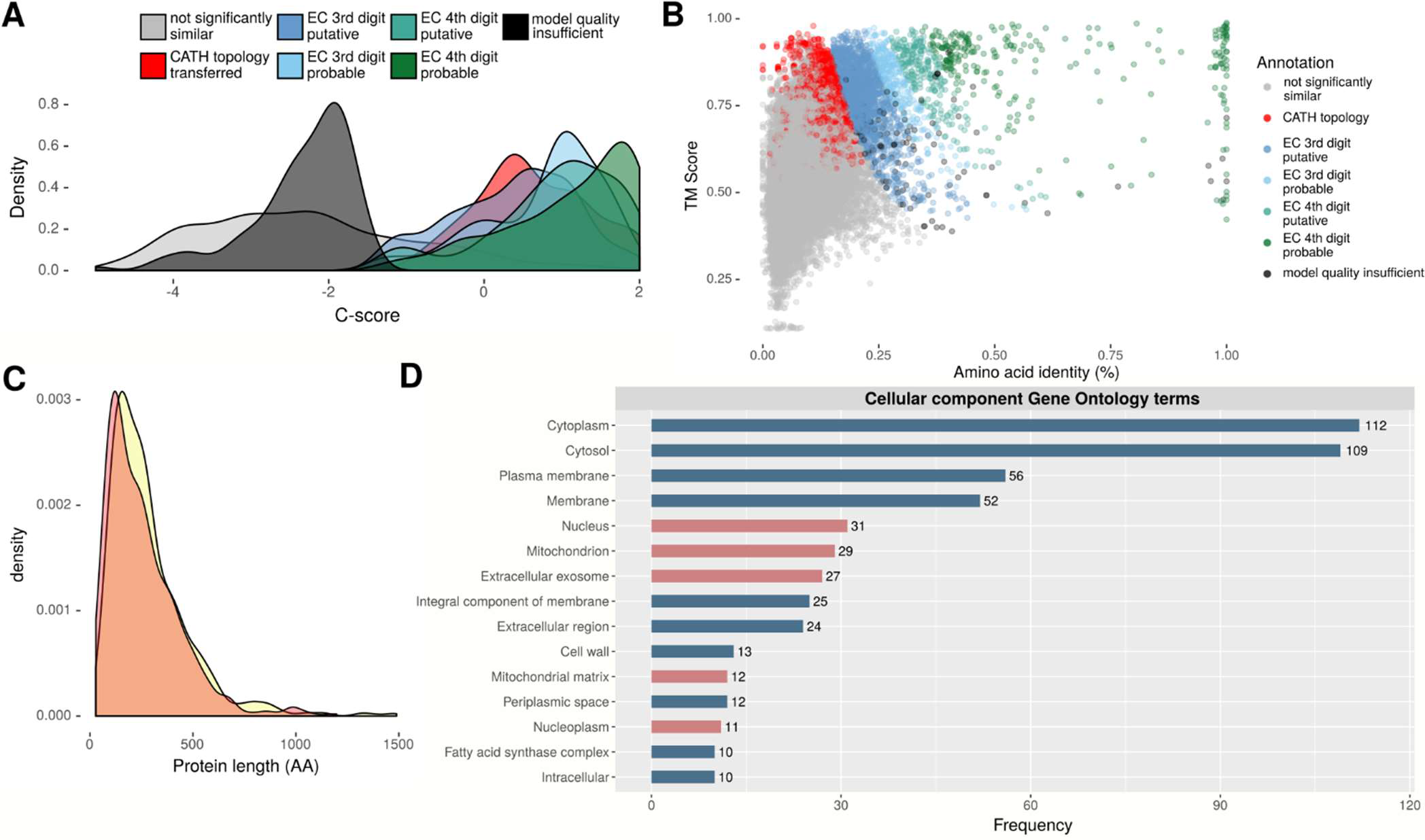
Relationships between model quality, annotation specificity, and protein features. **(A) Distribution of model quality (C-score) for levels of annotation confidence and specificity.** For each of the 1,711 protein models, the top three most similar PDB entries are included. Despite not explicitly accounting for differential degrees of error in C-score across proteins modeled, annotations were transferred higher quality models were provided EC thresholds correlated positively to model quality. **(B) Annotation classification as a function of amino acid identity and TM-score.** Each dot represents a relationship between a structure in PDB and a predicted structure of one of the 1,711 proteins for which structural models were able to be generated. Colors indicate which classification the relationship qualified for based on our inclusion criteria. Shades running from blue to green depict progressively stricter thresholds for EC inclusion tiers and confidence as determined by µ_geom_:precision relation (Fig. 1). Red dots indicate that similarity between the structure model/PDB structure was insufficient to qualify for an EC or GO annotation, but met the TM-scoreADJ criteria for transfer of CATH topology (blue and green points may have also qualified for CATH transfer). Black dots indicate that though the raw TM-score would qualify the relationship for transfer of EC/GO annotation, the model was of insufficient quality (C-score < -1.5), and the annotation was therefore not transferred. The sharp transitions in the distribution reflect the hierarchical implementation of inclusion criteria for functional annotations (EC number and GO terms). **(C) Protein length distributes similarly in models in high- and low-quality structural models.** Density in the distribution of protein length for model above (red) and below (yellow) the threshold for protein structure model quality (C-score > -1.5)(14, 21). **(D) Most frequently annotated Cellular Component Gene Ontology (GO) terms.** Cellular component ontology GO term describe proteins by the cellular location where they perform their function (28). GO terms were incorporated as described in the flow diagram of Fig. S3B. Only those terms with ten or more occurrences are plotted, and no attempt was made to collapse child ontologies into parents. GO terms implicating function specific to eukaryotic cellular components are shown in red.

## Supplemental material descriptions

**Dataset 1:** (A) **Under-annotated genes.** List of all under-annotated genes analyzed in the study, their reason for inclusion, and product annotation on TubercuList. (B) **Annotation summary.** A comprehensive annotation overview for all 1,725 under-annotated genes. For each under-annotated gene, annotations according to TubercuList, manual curation, structurally similar proteins from I-TASSER, CATH structural topologies, and Ligand Binding sites are listed, along with structural model quality score (C-score) and classification (poor, strong, no model). (C) **Novel manual annotations.** All genes with manually curated product annotations novel to TubercuList, UniProt, and Mtb Network Portal. Annotations from each source are enumerated, along with PubMed IDs for each publication from which manually curated annotations were derived. (D) **Training set genes.** The list of genes selected randomly among proteins with experimentally-characterized, reviewed functional annotations on from UniProt (n=200) and manually curated annotations with specific functions (n=163) that were used as the training set to determine the putative and probable thresholds for transferring functional annotations between structurally similar proteins. (E) **Manually annotated gene products.** All genes with manually curated annotations from the literature in this work, their product annotations, level of confidence, and the sources from which the annotations were curated. (F) **Still lacking annotation.** All genes lacking LBS, CATH, functionally informative literature notes, or product annotations. Contains information about alternative databases with functional information and structure-quality (C-score > -1.5) that may provide leads to functional characterization or hypothesis generation. (G) **Syntenic unknown clusters.** Clusters of ≥3 consecutive genes lacking annotation have their potential roles listed, based on gene neighborhood. (H) **Chromosomal bias.** All genes in TubercuList and their annotation status, functional categories, and chromosomal location data (e.g. lagging vs. leading, terminus-proximal, etc.) used for enrichment analyses to identify orientation and spatial biases for uncharacterized genes. (I) **EC assignments.** All Enzyme commission numbers assigned to under-annotated genes in the study. (J) **KEGG mappings.** Generic KEGG pathway and subsystems mappings for all fully specified EC numbers (all four digits) annotated in this study. Many EC numbers map to multiple pathways/subsystems.

**Dataset 2. Comparison of manually curated annotations to annotations in UniProt.** (A-D) Contains results of comparison of 283 manually annotated under-annotated genes to UniProt annotations to identify instances where:

1. UniProt annotations and annotations from this study were discordant.
2. UniProt annotations and annotations from this study were concordant.
3. UniProt annotation was more informative of the function of the under-annotated genes.
4. Annotation from this study was more informative of the function of the under-annotated genes or
5. UniProt does not have a function annotation for the under-annotated genes.

We then distinguished genes annotated solely as antigens from those with other annotations to identify instances where the under-annotated genes had non-antigenic other functions associated with it in UniProt or the manual curation. (E) **Comparison of annotations manual annotations novel to UniProt to Mtb Network Portal, PATRIC, and RefSeq.** Results of comparing 173 under-annotated genes with manually annotated products absent from UniProt to the Mtb Network Portal, PATRIC, and RefSeq databases. Genes novelty are indicated by cell color. <u>Yellow</u>: Genes given a novel annotation in one or more databases where other databases lacked or had ambiguous annotation. For example, Rv0394c is annotated as Hyaluronidase/Chondrosulfatase only in our manual annotation; it is annotated ambiguously (“membrane protein” or “hypothetical protein”) in other databases, so only the annotation in the “Our manual annotation” column is highlighted. <u>Magenta</u>: Genes given an annotation more informative or of greater confidence annotation compared to other analyzed databases. For example, Rv3632 is annotated as a flippase in our manual annotation; which is more specific than annotations of “membrane protein” or “cell division protein” in other databases. <u>Lime Green</u>: Gene annotations that matched between our manual annotation and other database annotations. <u>Cyan</u>: Gene annotations that were distinct in our manual annotation from one or more annotations in other databases such that it is possible our annotation and the other annotations could both describe the gene’s functional product. For example, Rv0020c is annotated as a Probable Cell wall synthesis protein with FHA domain, FhaA in our database and Mtb Network portal’s database, but these annotations differ from PATRIC’s annexin VII annotation, so the annotations in all three of these columns are highlighted. Empty rows indicate that a database did not have an annotation for the protein. This sheet can be converted to html format using Microsoft Excel/Open office/Libreoffice or an online file conversion tool for ease of programmatically reproducing counts. PATRIC comments are included in the analysis for comparison, but mostly contain putative annotations from Phyre2 ab initio prediction software, which are considered weaker than the experimental evidence found by manual curation.

**Dataset 3. Structure-based annotations.** (A) **Strong protein-PDB matches.** All matches between predicted 3D structure of proteins encoded by under-annotated genes and PDB structure entries. All matches that either exceeded the geometric mean cutoff for inclusion in annotation as “putative” (> 50% observed probability of correct annotation), or with an adjusted TM-score > 0.52 are included. PDB IDs, macromolecular names, GO terms of the PDB match, are included, as well as statistic of model quality and match strength. (B) **Top PDB matches.** For all proteins with a model generated, the top 3 PDB entry matches are listed, irrespective of similarity metrics. Similarity metrics are listed, along with information about the matching PDB entry. (C) **CATH families.** All CATH families for all structure-PDB entry matches with adjusted TM-score greater than 0.52. The coordinates annotated with each CATH are listed, along with similarity metrics and model quality (C-score). (D) **Ligand Binding Sites.** All predicted Ligand Binding Sites for which C-scoreLBS ≥0.6. (E) Transport-like PDB hits. All matches according to criteria for transporters in methods are included. Information about the matching PDB template (E.C. number, GO terms, macromolecular name) are included, as well as similarity metrics between 3D structure models and PDB entries. (F) **Corroborated transport proteins.** The top two Transport-like protein hits (ranked by TM_ADJ_) that passed the structure-inferred quality control pipeline. Sheets G-K contain orthogonal validation and manual structure analysis quality assurance steps. (G) **HHpred EC concordance.** EC numbers for each enzymatic annotations from proteins with functions inferred by structure alone. Left columns are the structure-inferred EC digits, and the rightmost four are those that were corroborated by HHpred hits. (H) **HHpred results.** Parsed HHpred output for all proteins with non-enzymatic annotations, alongside their structure-inferred product name. (I) **Consensus non-EC HHpred.** Sheet containing final reconciled product name corroborations, refutations, and modifications based on two-reviewer consensus of match/mismatch between HHpred hits and structure-inferred product annotations. (J) **Ramachandran filtering.** Fraction of residues in most favored, additional allowed, generously allowed, and disallowed regions for the proteins uncorroborated by HHpred, and some fully corroborated as positive controls used to build the threshold to define inclusion criteria to go on to manual structural analysis. (K) **Manual structure inspection.** Notes and results of manual verification of structure quality, structure-to-function mapping, and conservation of key structural features and residues for HHpred-uncorroborated proteins whose model structure passed Ramachandran filtering. (L) **CATH topologies.** Most commonly transferred CATH topologies and their associated functions. (M) **TetR matches.** Genes with PDB matches with Tetracyline Repressor 2 (TetR) CATH topologies. Only the highest TM_ADJ_ is shown. Amino acid coordinates annotated with the CATH topology are indicated (AA coords), along with the PDB template, model quality (C-Score), Amino acid identity to the PDB match (Identity), and raw TM-score (TM-Score). (N) **Mycobacterial core genes.** Genes identified recently as part of the mycobacterial core genome(67) and annotated as hypothetical proteins (according to Table S3 of the recent work) are listed, along with their updated annotation and its source(s).

**Text S1.** Further details of approach, methods, and results.

**Supplemental annotation file.** Full machine-readable H37Rv annotation file in GFF format. Contains all annotations, notes, and evidence from the study

## CITATIONS

1. Poux S, Arighi CN, Magrane M, Bateman A, Wei C-H, Lu Z, Boutet E, Bye-A-Jee H, Famiglietti ML, Roechert B. On expert curation and scalability: UniProtKB/ Swiss-Prot as a case study.

2. Steenken W, Oatway WH, Petroff SA. 1934. BIOLOGICAL STUDIES OF THE TUBERCLE BACILLUS. J Exp Med 60.

3. Cole ST, Brosch R, Parkhill J, Garnier T, Churcher C, Harris D, Gordon S V., Eiglmeier K, Gas S, Barry CE, Tekaia F, Badcock K, Basham D, Brown D, Chillingworth T, Connor R, Davies R, Devlin K, Feltwell T, Gentles S, Hamlin N, Holroyd S, Hornsby T, Jagels K, Krogh A, McLean J, Moule S, Murphy L, Oliver K, Osborne J, Quail MA, Rajandream M-A, Rogers J, Rutter S, Seeger K, Skelton J, Squares R, Squares S, Sulston JE, Taylor K, Whitehead S, Barrell BG. 1998. Deciphering the biology of Mycobacterium tuberculosis from the complete genome sequence. Nature 393:537–544.

4. Camus J-C, Pryor MJ, Médigue C, Cole ST. 2002. Re-annotation of the genome sequence of Mycobacterium tuberculosis H37Rv. Microbiology 148:2967–2973.

5. Reddy TBK, Riley R, Wymore F, Montgomery P, DeCaprio D, Engels R, Gellesch M, Hubble J, Jen D, Jin H, Koehrsen M, Larson L, Mao M, Nitzberg M, Sisk P, Stolte C, Weiner B, White J, Zachariah ZK, Sherlock G, Galagan JE, Ball CA, Schoolnik GK. 2009. TB database: an integrated platform for tuberculosis research. Nucleic Acids Res 37:D499–D508.

6. Gillespie JJ, Wattam AR, Cammer SA, Gabbard JL, Shukla MP, Dalay O, Driscoll T, Hix D, Mane SP, Mao C, Nordberg EK, Scott M, Schulman JR, Snyder EE, Sullivan DE, Wang C, Warren A, Williams KP, Xue T, Seung Yoo H, Zhang C, Zhang Y, Will R, Kenyon RW, Sobral BW. 2011. PATRIC: the Comprehensive Bacterial Bioinformatics Resource with a Focus on Human Pathogenic Species ▿. Infect Immun 79:4286–4298.

7. Garcia BJ, Datta G, Davidson RM, Strong M. 2015. MycoBASE: expanding the functional annotation coverage of mycobacterial genomes. BMC Genomics 16:1102.

8. Schito M, Dolinger DL. 2015. A Collaborative Approach for “ReSeq-ing” Mycobacterium tuberculosis Drug Resistance: Convergence for Drug and Diagnostic Developers. EBioMedicine 2:1262–1265.

9. Ma S, Minch KJ, Rustad TR, Hobbs S, Zhou S-L, Sherman DR, Price ND. 2015. Integrated Modeling of Gene Regulatory and Metabolic Networks in Mycobacterium tuberculosis. PLoS Comput Biol 11:e1004543.

10. Faksri K, Tan JH, Chaiprasert A, Teo Y-Y, Ong RT-H. 2016. Bioinformatics tools and databases for whole genome sequence analysis of Mycobacterium tuberculosis. Infect Genet Evol 45:359–368.

11. Silva D, Almeida PE, Von Groll A, Martin A, Palomino JC. 2011. Efflux as a mechanism for drug resistance in Mycobacterium tuberculosis. FEMS Immunol Med Microbiol 63:1–9.

12. Theobald DL, Miller C. 2010. Membrane transport proteins: surprises in structural sameness. Nat Struct Mol Biol 17:2–3.

13. Drayman N, Glick Y, Ben-nun-Shaul O, Zer H, Zlotnick A, Gerber D, Schueler-Furman O, Oppenheim A. 2013. Pathogens Use Structural Mimicry of Native Host Ligands as a Mechanism for Host Receptor Engagement. Cell Host Microbe 14:63–73.

14. Yang J, Yan R, Roy A, Xu D, Poisson J, Zhang Y. 2015. The I-TASSER Suite: protein structure and function prediction. Nat Methods 12:7–8.

15. Rose PW, Prlić A, Altunkaya A, Bi C, Bradley AR, Christie CH, Costanzo L Di, Duarte JM, Dutta S, Feng Z, Green RK, Goodsell DS, Hudson B, Kalro T, Lowe R, Peisach E, Randle C, Rose AS, Shao C, Tao Y-P, Valasatava Y, Voigt M, Westbrook JD, Woo J, Yang H, Young JY, Zardecki C, Berman HM, Burley SK. 2017. The RCSB protein data bank: integrative view of protein, gene and 3D structural information. Nucleic Acids Res 45:D271–D281.

16. Moult J, Fidelis K, Kryshtafovych A, Rost B, Hubbard T, Tramontano A. 2007. Critical assessment of methods of protein structure prediction—Round VII. Proteins Struct Funct Bioinforma 69:3–9.

17. Kryshtafovych A, Krysko O, Daniluk P, Dmytriv Z, Fidelis K. 2009. Protein structure prediction center in CASP8. Proteins Struct Funct Bioinforma 77:5–9.

18. Moult J, Fidelis K, Kryshtafovych A, Tramontano A. 2011. Critical assessment of methods of protein structure prediction (CASP)—round IX. Proteins Struct Funct Bioinforma 79:1–5.

19. Huang YJ, Mao B, Aramini JM, Montelione GT. 2014. Assessment of template-based protein structure predictions in CASP10. Proteins Struct Funct Bioinforma 82:43–56.

20. Moult J, Fidelis K, Kryshtafovych A, Schwede T, Tramontano A. 2016. Critical assessment of methods of protein structure prediction: Progress and new directions in round XI. Proteins Struct Funct Bioinforma 84:4–14.

21. Xu J, Zhang Y. 2010. How significant is a protein structure similarity with TM-score = 0.5? Bioinformatics 26:889–895.

22. Zhang Y, Skolnick J. 2005. TM-align: a protein structure alignment algorithm based on the TM-score. Nucleic Acids Res 33:2302–9.

23. Zhang C, Freddolino PL, Zhang Y. 2017. COFACTOR: improved protein function prediction by combining structure, sequence and protein–protein interaction information. Nucleic Acids Res 45:W291–W299.

24. Ashburner M, Ball CA, Blake JA, Botstein D, Butler H, Cherry JM, Davis AP, Dolinski K, Dwight SS, Eppig JT, Harris MA, Hill DP, Issel-Tarver L, Kasarskis A, Lewis S, Matese JC, Richardson JE, Ringwald M, Rubin GM, Sherlock G. 2000. Gene Ontology: tool for the unification of biology. Nat Genet 25:25–29.

25. Webb EC, others. 1992. Enzyme nomenclature 1992. Recommendations of the Nomenclature Committee of the International Union of Biochemistry and Molecular Biology on the Nomenclature and Classification of Enzymes. Academic Press.

26. Yang J, Roy A, Zhang Y. 2013. BioLiP: a semi-manually curated database for biologically relevant ligand-protein interactions. Nucleic Acids Res 41:D1096–103.

27. Hill DP, Davis AP, Richardson JE, Corradi JP, Ringwald M, Eppig JT, Blake JA. 2001. PROGRAM DESCRIPTION: Strategies for Biological Annotation of Mammalian Systems: Implementing Gene Ontologies in Mouse Genome Informatics. Genomics 74:121–128.

28. The Gene Ontology Consortium, Gene Ontology Consortium, The Gene Ontology Consortium. 2017. Expansion of the Gene Ontology knowledgebase and resources. Nucleic Acids Res 45:D331–D338.

29. Ramakrishnan G, Ochoa-Montaño B, Raghavender US, Mudgal R, Joshi AG, Chandra NR, Sowdhamini R, Blundell TL, Srinivasan N. 2015. Enriching the annotation of Mycobacterium tuberculosis H37Rv proteome using remote homology detection approaches: Insights into structure and function. Tuberculosis 95:14–25.

30. Functional assignment of Mycobacterium tuberculosis proteome revealed by genome-scale fold-recognition.

31. Al-Khafaji ZM. 2013. In Silico Investigation of Rv Hypothetical Proteins of Virulent Strain Mycobacterium tuberculosis H37Rv. ndian J Pharm Biol Res 1:81–88.

32. Doerks T, Noort V van, Minguez P, Bork P. 2012. Annotation of the *M. tuberculosis* Hypothetical Orfeome: Adding Functional Information to More than Half of the Uncharacterized Proteins. PLoS One 7:e34302.

33. Mazandu GK, Mulder NJ. 2012. Using the underlying biological organization of the Mycobacterium tuberculosis functional network for protein function prediction. Infect Genet Evol 12:922–932.

34. Sangrador-Vegas A, Mitchell AL, Chang H-Y, Yong S-Y, Finn RD. 2016. GO annotation in InterPro: why stability does not indicate accuracy in a sea of changing annotations. Database 2016:baw027.

35. Liberal R, Pinney JW. 2013. Simple topological properties predict functional misannotations in a metabolic network. Bioinformatics 29.

36. Tatusova T, Ciufo S, Fedorov B, O’Neill K, Tolstoy I. 2014. RefSeq microbial genomes database: new representation and annotation strategy. Nucleic Acids Res 42:D553–D559.

37. Sultana R, Vemula MH, Banerjee S, Guruprasad L. 2013. The PE16 (Rv1430) of Mycobacterium tuberculosis Is an Esterase Belonging to Serine Hydrolase Superfamily of Proteins. PLoS One https://doi.org/10.1371/journal.pone.0055320.

38. Padhi A, Naik SK, Sengupta S, Ganguli G, Sonawane A. 2016. Expression of Mycobacterium tuberculosis NLPC/p60 family protein Rv0024 induce biofilm formation and resistance against cell wall acting anti-tuberculosis drugs in Mycobacterium smegmatis. Microbes Infect 18:224–236.

39. Kateete DP, Katabazi FA, Okeng A, Okee M, Musinguzi C, Asiimwe BB, Kyobe S, Asiimwe J, Boom WH, Joloba ML. 2012. Rhomboids of Mycobacteria: Characterization Using an aarA Mutant of Providencia stuartii and Gene Deletion in Mycobacterium smegmatis. PLoS One https://doi.org/10.1371/journal.pone.0045741.

40. Nambi S, Long JE, Mishra BB, Baker R, Murphy KC, Olive AJ, Nguyen HP, Shaffer SA, Sassetti CM. 2015. The Oxidative Stress Network of Mycobacterium tuberculosis Reveals Coordination between Radical Detoxification Systems. Cell Host Microbe https://doi.org/10.1016/j.chom.2015.05.008.

41. Flentie K, Garner AL, Stallings CL. 2016. Mycobacterium tuberculosis transcription machinery: Ready to respond to host attacks. J Bacteriol.

42. Primm TP, Andersen SJ, Mizrahi V, Avarbock D, Rubin H, Barry Iii CE. 2000. The Stringent Response of Mycobacterium tuberculosis Is Required for Long-Term Survival. J Bacteriol 182:4889–4898.

43. Söding J, Biegert A, Lupas AN. 2005. The HHpred interactive server for protein homology detection and structure prediction. Nucleic Acids Res 33:W244.

44. Bhattacharya A, Tejero R, Montelione GT. 2007. Evaluating protein structures determined by structural genomics consortia. Proteins Struct Funct Genet 66:778–795.

45. Laskowski RA, MacArthur MW, Moss DS, Thornton JM. 1993. PROCHECK: a program to check the stereochemical quality of protein structures. J Appl Crystallogr 26:283–291.

46. He YX, Huang L, Xue Y, Fei X, Teng Y Bin, Rubin-Pitel SB, Zhao H, Zhou CZ. 2010. Crystal structure and computational analyses provide insights into the catalytic mechanism of 2,4-diacetylphloroglucinol hydrolase PhlG from Pseudomonas fluorescens. J Biol Chem 285:4603–4611.

47. Han J, Zhang S, Jia W, Zhang Z, Wang Y, He Y. 2019. Discovery and structural analysis of a phloretin hydrolase from the opportunistic human pathogen *Mycobacterium abscessus*. FEBS J 286:1959–1971.

48. Brzostek A, Pawelczyk J, Rumijowska-Galewicz A, Dziadek B, Dziadek J. 2009. Mycobacterium tuberculosis is able to accumulate and utilize cholesterol. J Bacteriol 191:6584–91.

49. Lack NA, Yam KC, Lowe EE, Horsman GP, Owen RL, Sim E, Eltis LD. 2010. Characterization of a carbon-carbon hydrolase from Mycobacterium tuberculosis involved in cholesterol metabolism. J Biol Chem 285:434–443.

50. Wilburn KM, Fieweger RA, VanderVen BC. 2018. Cholesterol and fatty acids grease the wheels of Mycobacterium tuberculosis pathogenesis. Pathog Dis. Oxford Academic.

51. Wang R, Yin YJ, Wang F, Li M, Feng J, Zhang HM, Zhang JP, Liu SJ, Chang WR. 2007. Crystal structures and site-directed mutagenesis of a mycothiol-dependent enzyme reveal a novel folding and molecular basis for mycothiol-mediated maleylpyruvate isomerization. J Biol Chem 282:16288–16294.

52. Newton GL, Leung SS, Wakabayashi JI, Rawat M, Fahey RC. 2011. The DinB Superfamily Includes Novel Mycothiol, Bacillithiol, and Glutathione S-Transferases. Biochemistry 50:10751–10760.

53. Liu TT, Zhou NY. 2012. Novel L-cysteine-dependent maleylpyruvate isomerase in the gentisate pathway of Paenibacillus sp. strain NyZ101. J Bacteriol 194:3987–3994.

54. Slama N, Jamet S, Frigui W, Pawlik A, Bottai D, Laval F, Constant P, Lemassu A, Cam K, Daffé M, Brosch R, Eynard N, Quémard A. 2015. The changes in mycolic acid structures caused by hadC mutation have a dramatic effect on the virulence of Mycobacterium tuberculosis. Mol Microbiol 99:n/a-n/a.

55. Singh A, Crossman DK, Mai D, Guidry L, Voskuil MI, Renfrow MB, Steyn AJC. 2009. Mycobacterium tuberculosis WhiB3 Maintains Redox Homeostasis by Regulating Virulence Lipid Anabolism to Modulate Macrophage Response. PLoS Pathog 5:e1000545.

56. Olivella M, Gonzalez A, Pardo L, Deupi X. 2013. Relation between sequence and structure in membrane proteins. Bioinformatics 29:1589–1592.

57. Shi W, Chen J, Zhang S, Zhang W, Zhang Y. 2018. Identification of novel mutations in LprG (rv1411c), rv0521, rv3630, rv0010c, ppsC, and cyp128 associated with pyrazinoic acid/pyrazinamide resistance in mycobacterium tuberculosis. Antimicrob Agents Chemother 62.

58. Costa P, Amaro A, Ferreira AS, Machado D, Albuquerque T, Couto I, Botelho A, Viveiros M, Inácio J. 2014. Rapid identification of veterinary-relevant Mycobacterium tuberculosis complex species using 16S rDNA, IS6110 and Regions of Difference-targeted dual-labelled hydrolysis probes. J Microbiol Methods 107:13–22.

59. Strong EJ, Jurcic Smith KL, Saini NK, Ng TW, Porcelli SA, Lee S. 2020. Identification of autophagy-inhibiting factors of mycobacterium tuberculosis by high-throughput loss-of-function screening. Infect Immun 88.

60. Kania E, Pająk B, O’Prey J, Sierra Gonzalez P, Litwiniuk A, Urbańska K, Ryan KM, Orzechowski A. 2017. Verapamil treatment induces cytoprotective autophagy by modulating cellular metabolism. FEBS J 284:1370–1387.

61. Abramovitch RB, Rohde KH, Hsu F-F, Russell DG. 2011. {aprABC}: {A} {Mycobacterium} tuberculosis complex-specific locus that modulates {pH}-driven adaptation to the macrophage phagosome. Mol Microbiol 80:678–694.

62. Agranoff D, Krishna S. 2004. Metal ion transport and regulation in {Mycobacterium} tuberculosis. Front Biosci A J Virtual Libr 9:2996–3006.

63. Rodrigues L, Cravo P, Viveiros M. 2020. Efflux pump inhibitors as a promising adjunct therapy against drug resistant tuberculosis: a new strategy to revisit mycobacterial targets and repurpose old drugs. Expert Rev Anti Infect Ther. Taylor and Francis Ltd.

64. Lin Y, Dong Y, Gao Y, Shi R, Li Y, Zhou X, Liu W, Li G, Qi Y, Wu Y. 2021. Identification of CTL Epitopes on Efflux Pumps of the ATP-Binding Cassette and the Major Facilitator Superfamily of Mycobacterium tuberculosis. J Immunol Res 2021:1–13.

65. Moult J, Melamud E. 2000. From fold to function. Curr Opin Struct Biol 10:384–389.

66. Evans JC, Mizrahi V. 2015. The application of tetracyclineregulated gene expression systems in the validation of novel drug targets in Mycobacterium tuberculosis. Front Microbiol 6.

67. Judd JA, Canestrari J, Clark R, Joseph A, Lapierre P, Lasek-Nesselquist E, Mir M, Palumbo M, Smith C, Stone M, Upadhyay A, Wirth SE, Dedrick RM, Meier CG, Russell DA, Dills A, Dove E, Kester J, Wolf ID, Zhu J, Rubin ER, Fortune S, Hatfull GF, Gray TA, Wade JT, Derbyshire KM. 2021. A mycobacterial systems resource for the research community. MBio 12:1–15.

68. Lato DF, Golding GB. 2020. Spatial Patterns of Gene Expression in Bacterial Genomes. J Mol Evol 88:510–520.

69. Eisen JA, Heidelberg JF, White O, Salzberg SL. 2000. Evidence for symmetric chromosomal inversions around the replication origin in bacteria. Genome Biol 1:research0011.1.

70. Repar J, Warnecke T. 2017. Non-Random Inversion Landscapes in Prokaryotic Genomes Are Shaped by Heterogeneous Selection Pressures. Mol Biol Evol 34:1902–1911.

71. Galperin MY, Kristensen DM, Makarova KS, Wolf YI, Koonin E V. 2019. Microbial genome analysis: the COG approach. Brief Bioinform 20:1063–1070.

72. Merrikh CN, Merrikh H. 2018. Gene inversion potentiates bacterial evolvability and virulence. Nat Commun 9:4662.

73. Onwueme KC, Vos CJ, Zurita J, Ferreras JA, Quadri LEN. 2005. The dimycocerosate ester polyketide virulence factors of mycobacteria. Prog Lipid Res 44:259–302.

74. Lee JS, Krause R, Schreiber J, Mollenkopf H-J, Kowall J, Stein R, Jeon B-Y, Kwak J-Y, Song M-K, Patron JP, Jorg S, Roh K, Cho S-N, Kaufmann SHE. 2008. Mutation in the Transcriptional Regulator PhoP Contributes to Avirulence of Mycobacterium tuberculosis H37Ra Strain. Cell Host Microbe 3:97–103.

75. Elghraoui A, Modlin SJ, Valafar F. 2017. SMRT genome assembly corrects reference errors, resolving the genetic basis of virulence in Mycobacterium tuberculosis. BMC Genomics 18:302.

76. Passemar C, Arbués A, Malaga W, Mercier I, Moreau F, Lepourry L, Neyrolles O, Guilhot C, Astarie-Dequeker C. 2014. Multiple deletions in the polyketide synthase gene repertoire of Mycobacterium tuberculosis reveal functional overlap of cell envelope lipids in host–pathogen interactions. Cell Microbiol 16:195–213.

77. Minnikin DE, Dobson G, Hutchinson IG. 1983. Characterization of phthiocerol dimycocerosates from Mycobacterium tuberculosis. Biochim Biophys Acta - Lipids Lipid Metab 753:445–449.

78. Mann FM, Xu M, Davenport EK, Peters RJ. 2012. Functional characterization and evolution of the isotuberculosinol operon in Mycobacterium tuberculosis and related Mycobacteria. Front Microbiol 3.

79. Mann FM, Prisic S, Hu H, Xu M, Coates RM, Peters RJ. 2009. Characterization and Inhibition of a Class II Diterpene Cyclase from Mycobacterium tuberculosis. J Biol Chem 284:23574–23579.

80. Mann FM, Peters RJ. 2012. Isotuberculosinol: the unusual case of an immunomodulatory diterpenoid from Mycobacterium tuberculosis. Medchemcomm 3:899–904.

81. Sieniawska E, Swatko-Ossor M, Sawicki R, Skalicka-Woźniak K, Ginalska G. 2017. Natural Terpenes Influence the Activity of Antibiotics against Isolated Mycobacterium tuberculosis. Med Princ Pract Int J Kuwait Univ Heal Sci Cent 26:108–112.

82. Layre E, Lee HJ, Young DC, Martinot AJ, Buter J, Minnaard AJ, Annand JW, Fortune SM, Snider BB, Matsunaga I, Rubin EJ, Alber T, Moody DB. 2014. Molecular profiling of Mycobacterium tuberculosis identifies tuberculosinyl nucleoside products of the virulence-associated enzyme Rv3378c. Proc Natl Acad Sci 111:2978–2983.

83. Bellerose MM, Proulx MK, Smith CM, Baker RE, Ioerger TR, Sassetti CM. 2020. Distinct Bacterial Pathways Influence the Efficacy of Antibiotics against Mycobacterium tuberculosis. mSystems 5.

84. Jansen RS, Mandyoli L, Hughes R, Wakabayashi S, Pinkham JT, Selbach B, Guinn KM, Rubin EJ, Sacchettini JC, Rhee KY. 2020. Aspartate aminotransferase Rv3722c governs aspartate-dependent nitrogen metabolism in Mycobacterium tuberculosis. Nat Commun 11:1–13.

85. Zhang L, Hendrickson RC, Meikle V, Lefkowitz EJ, Ioerger TR, Niederweis M. 2020. Comprehensive analysis of iron utilization by Mycobacterium tuberculosis. PLoS Pathog 16:e1008337.

86. Scorpio A, Zhang Y. 1996. Mutations in pncA, a gene encoding pyrazinamidase/nicotinamidase, cause resistance to the antituberculous drug pyrazinamide in tubercle bacillus. Nat Med 2:662–667.

87. Zhang Y, Wade MM, Scorpio A, Zhang H, Sun Z. 2003. Mode of action of pyrazinamide: Disruption of Mycobacterium tuberculosis membrane transport and energetics by pyrazinoic acid. J Antimicrob Chemother 52:790–795.

88. Zimhony O, Cox JS, Welch JT, Vilchèze C, Jacobs WR. 2000. Pyrazinamide inhibits the eukaryotic-like fatty acid synthetase I (FASI) of Mycobacterium tuberculosis. Nat Med 6:1043–1047.

89. Shi W, Zhang X, Jiang X, Yuan H, Lee JS, Barry CE, Wang H, Zhang W, Zhang Y. 2011. Pyrazinamide inhibits trans-translation in Mycobacterium tuberculosis. Science (80-) 333:1630–1632.

90. Boshoff HI, Mizrahi V, Iii CEB, Barry CE. 2002. Effects of Pyrazinamide on Fatty Acid Synthesis by Whole Mycobacterial Cells and Purified Fatty Acid Synthase I. J Bacteriol 184:2167–2172.

91. Peterson ND, Rosen BC, Dillon NA, Baughn AD. 2015. Uncoupling environmental pH and intrabacterial acidification from pyrazinamide susceptibility in Mycobacterium tuberculosis. Antimicrob Agents Chemother 59:7320–7326.

92. Dillon NA, Peterson ND, Feaga HA, Keiler KC, Baughn AD. 2017. Anti-tubercular Activity of Pyrazinamide is Independent of trans-Translation and RpsA. Sci Rep 7:6135.

93. Gopal P, Yee M, Sarathy JPJ, Liang Low J, Sarathy JPJ, Kaya F, Dartois VV, Gengenbacher M, Dick T, Low JL, Sarathy JPJ, Kaya F, Dartois VV, Gengenbacher M, Dick T. 2016. Pyrazinamide resistance is caused by two distinct mechanisms: Prevention of coenzyme a depletion and loss of virulence factor synthesis. ACS Infect Dis 2:616–626.

94. Anthony RM, den Hertog AL, van Soolingen D. 2018. ‘Happy the man, who, studying nature’s laws, Thro’’ known effects can trace the secret cause.’† Do we have enough pieces to solve the pyrazinamide puzzle?’ J Antimicrob Chemother 73:1750– 1754.

95. Dillon NA, Peterson ND, Rosen BCBC, Baughn AD. 2014. Pantothenate and pantetheine antagonize the antitubercular activity of pyrazinamide. Antimicrob Agents Chemother 58:7258–7263.

96. Zhang Y, Mitchison D. 2003. The curious characteristics of pyrazinamide: A review. Int J Tuberc Lung Dis 7:6–21.

97. Swan MK, Hansen T, Schönheit P, Davies C. 2004. Structural basis for phosphomannose isomerase activity in phosphoglucose isomerase from Pyrobaculum aerophilum: A subtle difference between distantly related enzymes. Biochemistry 43:14088–14095.

98. Mouilleron S, Badet-Denisot MA, Golinelli-Pimpaneau B. 2006. Glutamine binding opens the ammonia channel and activates glucosamine-6P synthase. J Biol Chem 281:4404–4412.

99. Bateman A. 1999. The SIS domain: A phosphosugar-binding domain. Trends Biochem Sci 24:94–95.

100. Keiser TL, Schlesinger LS, King S, Munson R, Seveau S. 2014. Biosynthesis of mannose-containing cell wall components important in Mycobacterium.

101. Schnoes AM, Brown SD, Dodevski I, Babbitt PC. 2009. Annotation Error in Public Databases: Misannotation of Molecular Function in Enzyme Superfamilies. PLoS Comput Biol 5.

102. Ioerger TR, Feng Y, Ganesula K, Chen X, Dobos KM, Fortune S, Jacobs WR, Mizrahi V, Parish T, Rubin E, Sassetti C, Sacchettini JC. 2010. Variation among Genome Sequences of H37Rv Strains of Mycobacterium tuberculosis from Multiple Laboratories. J Bacteriol 192:3645–3653.

103. Shell SS, Wang J, Lapierre P, Mir M, Chase MR, Pyle MM, Gawande R, Ahmad R, Sarracino DA, Ioerger TR, Fortune SM, Derbyshire KM, Wade JT, Gray TA. 2015. Leaderless Transcripts and Small Proteins Are Common Features of the Mycobacterial Translational Landscape. PLoS Genet 11.

104. Nagano N, Orengo CA, Thornton JM. 2002. One fold with many functions: the evolutionary relationships between TIM barrel families based on their sequences, structures and functions. J Mol Biol 321:741–765.

105. Murphy GS, Greisman JB, Hecht MH. 2016. De Novo Proteins with Life-Sustaining Functions Are Structurally Dynamic. J Mol Biol 428:399–411.

106. Miskei M, Gregus A, Sharma R, Duro N, Zsolyomi F, Fuxreiter M. 2017. Fuzziness enables context dependence of protein interactions. FEBS Lett 591:2682–2695.

107. Morcos F, Pagnani A, Lunt B, Bertolino A, Marks DS, Sander C, Zecchina R, Onuchic JN, Hwa T, Weigt M. 2011. Direct-coupling analysis of residue coevolution captures native contacts across many protein families. Proc Natl Acad Sci U S A 108:E1293–E1301.

108. Lockless SW, Ranganathan R. 1999. Evolutionarily conserved pathways of energetic connectivity in protein families. Science (80-) 286:295–299.

109. Neuwald AF. 2014. A Bayesian Sampler for Optimization of Protein Domain Hierarchies. J Comput Biol 21:269–286.

110. Neuwald AF, Aravind L, Altschul SF. 2018. Inferring joint sequence-structural determinants of protein functional specificity. Elife 7.

111. PDB RCSB. PDB Data Distribution by Resolution.

112. Jensen MR, Zweckstetter M, Huang J, Blackledge M. 2014. Exploring Free-Energy Landscapes of Intrinsically Disordered Proteins at Atomic Resolution Using NMR Spectroscopy. Chem Rev 114:6632–6660.

113. Fishbein S, van Wyk N, Warren RM, Sampson SL. 2015. Phylogeny to function: PE/PPE protein evolution and impact on Mycobacterium tuberculosis pathogenicity. Mol Microbiol 96:901–916.

114. McEvoy CREE, Cloete R, Müller B, Schürch AC, Helden PD van, Gagneux S, Warren RM, Pittius NCG van, van Helden PD, Gagneux S, Warren RM, Gey van Pittius NC. 2012. Comparative Analysis of Mycobacterium tuberculosis pe and ppe Genes Reveals High Sequence Variation and an Apparent Absence of Selective Constraints. PLoS One 7:e30593.

115. Zhang Y, Skolnick J. 2004. Scoring function for automated assessment of protein structure template quality. Proteins Struct Funct Bioinforma 57:702–710.

116. UniProt Consortium. 2017. UniProt: the universal protein knowledgebase. Nucleic Acids Res 45:D158–D169.

117. Roy A, Yang J, Zhang Y. 2012. COFACTOR: an accurate comparative algorithm for structure-based protein function annotation. Nucleic Acids Res 40:W471–W477.

118. R Development Core Team. 2018. A Language and Environment for Statistical Computing. R Found Stat Comput. Vienna, Austria.

119. Karp PD, Billington R, Caspi R, Fulcher CA, Latendresse M, Kothari A, Keseler IM, Krummenacker M, Midford PE, Ong Q, Ong WK, Paley SM, Subhraveti P. 2017. The BioCyc collection of microbial genomes and metabolic pathways. Brief Bioinform 28:1–6.

120. Kanehisa M, Furumichi M, Tanabe M, Sato Y, Morishima K. KEGG: new perspectives on genomes, pathways, diseases and drugs. Nucleic Acids Res 45.

121. Laskowski RA, MacArthur MW, Thornton JM. 2012. *PROCHECK*: validation of protein-structure coordinates, p. 684–687. *In*. American Cancer Society.

122. Rubin EJ, Akerley BJ, Novik VN, Lampe DJ, Husson RN, Mekalanos JJ. 1999. In vivo transposition of mariner-based elements in enteric bacteria and mycobacteria. Proc Natl Acad Sci 96:1645–1650.

123. Lee S, Kriakov J, Vilcheze C, Dai Z, Hatfull GF, Jacobs WR, J.F. C, J. M, D. M, N.R. P, W. B, V. K, J. K, L. K, S. B, J. K, J.G. L, W.R. J, R.W. H, G.F. H. 2004. Bxz1, a new generalized transducing phage for mycobacteria. FEMS Microbiol Lett 241:271–276.

124. Baughn AD, Deng J, Vilcheze C, Riestra A, Welch JT, Jacobs WR, Zimhony O, Vilchèze C, Riestra A, Welch JT, Jacobs WR, Zimhony O. 2010. Mutually exclusive genotypes for pyrazinamide and 5-chloropyrazinamide resistance reveal a potential resistance-proofing strategy. Antimicrob Agents Chemother 54:5323–5328.

125. Lew JM, Kapopoulou A, Jones LM, Cole ST. 2011. TubercuList--10 years after. Tuberculosis (Edinb) 91:1–7.

126. Chen J, Guo M, Wang X,Liu B. A comprehensive review and comparison of different computational methods for protein remote homology detection. Brief Bioinform https://doi.org/10.1093/bib/bbw108.

127. Cortes T, Schubert OTT, Rose G, Arnvig KBB, Comas I, Aebersold R, Young DBB. 2013. Genome-wide Mapping of Transcriptional Start Sites Defines an Extensive Leaderless Transcriptome in Mycobacterium tuberculosis. Cell Rep 5:1121– 1131.

128. Shell SS, Wang J, Lapierre P, Mir M, Chase MR, Pyle MM, Gawande R, Ahmad R, Sarracino DA, Ioerger TR, Fortune SM, Derbyshire KM, Wade JT, Gray TA. 2015. Leaderless Transcripts and Small Proteins Are Common Features of the Mycobacterial Translational Landscape. PLoS Genet 11.

129. Teplyakov A, Obmolova G, Badet-Denisot MA, Badet B, Polikarpov I. 1998. Involvement of the C terminus in intramolecular nitrogen channeling in glucosamine 6-phosphate synthase: Evidence from a 1.6 Å crystal structure of the isomerase domain. Structure 6:1047–1055.

130. Salah Ud-Din A, Tikhomirova A, Roujeinikova A. 2016. Structure and Functional Diversity of GCN5-Related N-Acetyltransferases (GNAT). Int J Mol Sci 17:1018.

131. Wybenga-Groot LE, Draker K ann, Wright GD, Berghuis AM. 1999. Crystal structure of an aminoglycoside 6’-N-acetyltransferase: Defining the GCN5-related N-acetyltransferase superfamily fold. Structure 7:497–507.

132. Reddy TBK, Riley R, Wymore F, Montgomery P, DeCaprio D, Engels R, Gellesch M, Hubble J, Jen D, Jin H, Koehrsen M, Larson L, Mao M, Nitzberg M, Sisk P, Stolte C, Weiner B, White J, Zachariah ZK, Sherlock G, Galagan JE, Ball CA, Schoolnik GK. 2009. TB database: an integrated platform for tuberculosis research. Nucleic Acids Res 37:D499–D508.

